# Barley powdery mildew invasion coincides with the dynamic accumulation of leaf apoplastic extracellular vesicles that are associated with host stress response proteins

**DOI:** 10.1101/2024.04.17.589905

**Authors:** Hannah Thieron, Pietro Spanu, Miriam Buhl, Charlotte Hülsmann, Charlotte Kummer, Fatih Demir, Pitter Huesgen, Ralph Panstruga

**Affiliations:** Unit for Plant Molecular Cell Biology, Institute for Biology I, RWTH Aachen University, 52074 Aachen, Germany; Imperial College, Department of Life Sciences, Imperial College Road, London, SW7 2AZ, London, U.K; Electron Microscopy Facility, Institute of Pathology, RWTH Aachen University Hospital, 52074 Aachen, Germany; Molecular and Biological Systems Analytics, ZEA-3 Analytics, Forschungszentrum Jülich, 52428 Jülich, Germany; Biochemistry and Functional Proteomics, Institute for Biology II, University of Freiburg, 79104 Freiburg, Germany; CIBSS - Centre for Integrative Biological Signaling Studies, University of Freiburg, 79104 Freiburg, Germany; Current address: Department of Biomedicine, Aarhus University, 8000 Aarhus C, Denmark

## Abstract

The mutual exchange of extracellular vesicles across kingdom borders is a feature of many plant-microbe interactions. The occurrence and cargos of extracellular vesicles has been studied in several instances, but their dynamics in the course of infection have remained elusive. Here we used two different procedures, differential high-speed centrifugation and polymer-based enrichment, to collect extracellular vesicles from the apoplastic wash fluid of barley (*Hordeum vulgare*) leaves challenged by its fungal powdery mildew pathogen, *Blumeria hordei*. Both methods yielded extracellular vesicles of similar quality and morphological characteristics, though the polymer approach was associated with higher reproducibility. We noted that extracellular vesicles derived from the apoplastic wash fluid constitute polydisperse populations that are selectively responsive to leaf infection by *B. hordei*. Extracellular vesicles of ∼100 nm – 300 nm diameter became progressively more abundant, in particular from 72 hours post inoculation onwards, resulting in a major peak late during fungal infection. Vesicles of ∼300 nm – 500 nm showed similar accumulation dynamics but reached much lower levels, suggesting they might constitute a separate population. Proteome analysis uncovered an enrichment of biotic stress response proteins associated with the extracellular vesicles. The barley t-SNARE protein Ror2, the ortholog of the PEN1 marker protein of extracellular vesicles in *Arabidopsis thaliana*, accumulates in extracellular vesicles during powdery mildew infection, hence also qualifying as a potential marker protein. Our study serves as a starting point for investigating the role of extracellular vesicles at different stages of plant-microbe interactions.

## Introduction

Extracellular vesicles (EVs) are evolutionarily conserved structures of ∼30-1,000 nm in size surrounded by a lipid bilayer and secreted by cells into the extracellular space (Colombo *et al*. 2014). During the last decades, their role as mediators of intra- and inter-organismal cell-cell communication has been established in various kingdoms of life, inducing physiologically relevant changes in recipient cells (Tkach & Théry 2016; Rybak & Robatzek 2019; U Stotz *et al*. 2022). Particularly well-explored in mammalian studies, their diverse cargo biomolecules, in various cell states and diseases underscore their potential for biomarker development and novel therapeutics (Herrmann *et al*. 2021; Stranford & Leonard 2017).

The presence of EVs in plants and their potential as mediators of cell-cell communication remained unremarked for over 50 years after their first description (Aist & Williams 1971; Halperin & Jensen 1967; Shaw & Manocha 1965). However, this topic has gained increasing attention in plant biology in recent years. EVs have been observed in apoplastic wash fluids (AWF) derived from various plant parts — leaves, roots, imbibing seeds, pollen, and fruits (Rutter & Innes 2017; Cai *et al*. 2018; Ju *et al*. 2013; Prado *et al*. 2014). In plant-microbe interactions, evidence is accumulating to suggest the involvement of EVs in the mutual manipulation of plant hosts and their colonizing microbes (Cai *et al*. 2021; Qiao *et al*. 2023). In such encounters, EVs were first discovered in *Arabidopsis thaliana* infected by the bacterial pathogen *Pseudomonas syringae* (Rutter & Innes 2017). Various types of cargo molecules, including nucleic acids (Ruf *et al*. 2022), proteins (He *et al*. 2021), and cell wall components (La Canal & Pinedo 2018; Bellis *et al*. 2022) have been found in association with EVs. Small RNAs of 18-25 nucleotides in size are frequently described as associated with EVs and are thought to mediate mutual cross-kingdom gene silencing in pathogenic (Cai *et al*. 2019) and mutualistic (Qiao *et al*. 2023) plant-microbe encounters. Apart from canonical small RNAs, circular RNAs (Zand Karimi *et al*. 2022), rRNA fragments (Kusch *et al*. 2023; Panstruga & Spanu 2024) and mRNAs (Kwon *et al*. 2021; Ruf *et al*. 2022; Wang *et al*. 2024) have been linked to these vesicles. Proteins associated with plant EVs are often enriched in stress- and immunity-related proteins, but also RNA-binding proteins possibly involved in RNA-loading onto EVs (Rutter & Innes 2017; Regente *et al*. 2017; He *et al*. 2021). Microbial EVs are generally thought to promote plant colonisation and were reported to be connected to modulating primary metabolism and virulence (Hill & Solomon 2020; Solé *et al*. 2015; Rutter *et al*. 2022). Despite the well-established association of different types of biomolecules with plant- and microbe-derived EVs, the precise localisation of these, inside or outside of EVs, remains in many cases unresolved and a matter of an ongoing debate (Nasfi & Kogel 2022; Zand Karimi *et al*. 2022; Cai *et al*. 2018).

EVs have diverse biogenesis pathways, which are best characterised in the animal/human field; they are broadly classified into apoptotic bodies from dying cells, microvesicles shedding off the plasma membrane, and exosomes derived from multivesicular bodies (Greening & Simpson 2018; Cocucci & Meldolesi 2015). In addition to these categories, at least two plant-specific EV types—exocyst-positive organelle-derived EVs (EXPO) and membrane tubules—have been proposed (Roth *et al*. 2019; Wang *et al*. 2010). The identification of marker proteins potentially discriminating EVs of different origin is pivotal for exploring EV functions. These markers for example, can, be used to track EV subpopulations by immunological or microscopy techniques (Bağcı *et al*. 2022). In Arabidopsis, the tetraspanin TET8 has been suggested as a marker for exosome-like EVs since it colocalises with the multivesicular body marker ARA6 (Cai *et al*. 2018). Other EV populations in Arabidopsis are identified by the patellin protein PATL1 and the syntaxin (t-SNARE protein) PEN1 (Rutter & Innes 2017).

Powdery mildew is a common disease of angiosperm plants. It is caused by obligate biotrophic ascomycete fungi that colonise above ground plant tissues (Glawe 2008). Accumulating evidence points to a cross-kingdom transfer of EVs in the course of plant-powdery mildew interactions. For example, upon infection of barley (*Hordeum vulgare*) and *Arabidopsis thaliana* with their compatible powdery mildew fungi, multivesicular bodies accumulate at plant-pathogen contact sites (An *et al*. 2006; Micali *et al*. 2011). The fusion of multivesicular bodies with the plasma membrane not only results in the direct release of molecules from the lumen into the extracellular space but also supposedly leads to the discharge of intraluminal vesicles, generating exosome-like EVs (Micali *et al*. 2011). Moreover, both the Arabidopsis EV marker PEN1 and its barley ortholog, Ror2, associate with multivesicular bodies and have been observed to accumulate extracellularly in pathogen-induced cell wall appositions (papillae) during both compatible and incompatible interactions (Bhat *et al*. 2005; Assaad *et al*. 2004; Böhlenius *et al*. 2010; Meyer *et al*. 2009). Finally, we and others gathered evidence, both *in silico* and *in vivo*, suggesting the natural exchange of sRNAs in the barley-*Blumeria hordei* pathosystem, potentially facilitated through EVs (Kusch *et al*. 2018; Kusch *et al*. 2023; Hunt *et al*. 2019). This notion is further corroborated by the earlier discovery of the phenomenon of “host-induced gene silencing” (HIGS), i.e. the suppression of pathogen transcripts in their natural environment by the expression of complementary double-stranded RNAs inside host cells, in the context of the barley-*B. hordei* interaction (Nowara *et al*. 2010).

In general, separation and characterisation of plant EVs follow procedures similar to those established in other organisms. In plants, leaf AWF typically serves as the source material, which differs from body fluids or cell culture media often used in other systems (Rutter & Innes 2017). Differential centrifugation remains the method of choice for EV enrichment in both plants and other types of organisms (Gardiner *et al*. 2016). However, the long handling times pose a severe disadvantage for this experimental route. Polymer-mediated precipitations, now often used in other organisms for EV isolation and functional studies, have not been employed in plants yet (Martínez-Greene *et al*. 2021). This isolation procedure relies on niches in polymer structures to capture EVs (Grunt *et al*. 2020). Irrespective of the first preparative step, usually a second purification step is conducted to reduce co-purifying contaminants or separate EV subtypes. The identified EV content depends on both the isolation procedure and the efforts to distinguish genuine EV cargo and from co-purifying contaminants (Clos-Sansalvador *et al*. 2022; Taylor & Shah 2015).

Here we explored further the role of EVs in the interaction between barley and its obligate biotrophic fungal pathogen, *B. hordei*. We undertook a comparative characterisation of EVs isolated using two distinct EV isolation protocols (differential ultracentrifugation and polymer-based enrichment (PME)) and characterised the obtained EVs regarding their morphology and abundance by transmission electron microscopy and nanoparticle tracking analysis (NTA). Through a time-course experiment spanning all developmental stages of the asexual *B. hordei* life cycle, we monitored the size and abundance of EVs and tracked the accumulation of the Arabidopsis EV marker ortholog Ror2. Complementing this comprehensive characterisation, we compiled an EV proteome catalogue and assessed potential marker proteins using a protease protection assay and size exclusion chromatography.

## Materials and Methods

### Plant and pathogen cultivation

Barley (*H. vulgare* L. cv. ”Margret“) was grown in So-Mi513 soil (HAWITA, Vechta, Germany) or Bio Topferde torffrei soil (HAWITA, Vechta, Germany) under a long day cycle (16 h light at 23 °C, 8 h darkness at 20 °C) at 60%-65% relative humidity and a light intensity of 105-120 μmol s^-1^ m^-2^. Seven-day-old barley plants were inoculated with *B. hordei* strain K1; the infected plants were kept under controlled conditions in a growth chamber with a long day cycle (12 h light at 20 °C, 12 h dark at 19 °C) at ca. 60% relative humidity and 100 μmol s^-1^ m^-2^ light intensity.

### AWF extraction and EV isolation

AWF was extracted from non-inoculated barely plants or after inoculation with *B. hordei* strain K1. Trays of both inoculated and non-inoculated plants were covered with lids and incubated prior to AWF extraction. Approximately 30 g of leaf fresh weight was collected and vacuum-infiltrated with potassium vesicle isolation buffer (Kusch *et al*. 2023). Excess buffer was carefully removed and the leaves were placed with cut ends down in 20-ml syringes. The syringes were inserted into 50-ml centrifuge tubes. AWF was collected by centrifugation at 400 *g* for 12 min at 4 °C. Cellular debris was removed by passing the AWF through a 0.45-µm syringe filter and further centrifugation at 10,000 *g* for 30 min at 4 °C. EVs were isolated from AWF by ultracentrifugation according to a recently published protocol (Rutter *et al*. 2017). The AWF was first centrifuged for 1 h at 40,000 g (4 °C) and the supernatant subsequently for 1 h at 100,000 *g* (4 °C) to collect two EV fractions, termed P40 and P100. Alternatively, a single EV sample was collected by PME using the “PME exosomes enrichment kit” (IST Innuscreen, Berlin, Germany) according to the manufacturer’s instructions. All EV pellets were dissolved in 20 mM Tris-HCl (pH 7.5). When indicated, EVs were further purified by passing through qEVoriginal/70 nm size exclusion chromatography columns (IZON, Lyon, France). EVs were either stored at 4 °C if morphology was analysed or snap-frozen and stored at -80 °C for other types of subsequent analysis.

### Trypan blue staining

Staining with trypan blue was performed as described before (Mulaosmanovic *et al*. 2020). The barley leaves were cleared in ethanol:acetic acid (3:1) for two days. The cleared leaves were then stained using 0.01% trypan blue in dH_2_O (w/v) for four hours. Leaves were washed and then stored in dH_2_O.

### Protein extraction and immunoblot analysis

Total protein extracts were prepared as described previously (Rutter & Innes 2017) by homogenising approximately 200 µl frozen leaf tissue in 400 µl protein extraction buffer (150 mM NaCl, 50 mM Tris pH 7.5, 0.1% (v/v)) and centrifuging for 10 min at 12,000 *g* (4 °C) to remove cell debris. Alternatively, total protein extraction was performed using phenol as described (Thomas *et al*. 2015). Proteins were denatured in 6x loading buffer (12% SDS, 47% glycerol, 60 mM Tris pH 6.8, 9% DTT (v/v)), by heating at 99 °C for 10 min and then subjected to sodium dodecyl sulfate-polyacrylamide gel electrophoresis. When indicated, protein gels were stained with silver as described (Chevallet *et al*. 2006) or Quick Coomassie Stain (Protein Ark, Rotherham, UK) according to the manufacturer’s instructions. Alternatively, proteins were transferred to a nitrocellulose membrane and used for immunodetection. Antibodies were purchased from Eurogentec (Liège, Belgium, custom-made polyclonal αRor2, (Collins *et al*. 2003), Agrisera (Vännäs, Sweden, αPsbA/D1 - AS05 084, αPR-1 - AS10 687, αRbcL - AS03 037, αCPN60A1 - AS12 2613), Dako A/S custom (Glostrup, Denmark, αPR- 17b, provided by Hans-Thordal Christensen lab) or Cell Signaling Technologies (Danvers, MA, USA, α-rabbit IgG-HRP, #7074). Chemiluminescence detection of antigen-antibody complexes was performed with SuperSignal^TM^ West Pico or Femto Western substrate (Thermo Fisher Scientific, Darmstadt, Germany). As a loading control, membranes were stained in Ponceau S (Applichem, Darmstadt, Germany) solution (0.05% (w/v) in 5% (v/v) acetic acid).

### Transmission electron microscopy

EVs in 0.2 M HEPES (pH 7.5) were allowed to adsorb on glow discharged formvar-carbon-coated nickel grids (Maxtaform, 200 mesh, Science Services GmbH, Munich, Germany) for 7 min. Negative staining was performed with 0.5% uranyl acetate (in aqua dest., Science Services GmbH, Munich, Germany) or 1% phosphotungstic acid (in aqua dest., Science Services GmbH, Munich, Germany). Grids were air-dried and imaged using a Hitachi HT7800 transmission electron microscope (Hitachi, Tokyo, Japan) operating at an acceleration voltage of 100 kV.

### NTA analysis

NTA analysis was performed with a NanoSight NS300 and NanoSight software version 3.2 (Malvern, Worcestershire, UK). EV samples were diluted with MilliQ H_2_O to a final volume of 1 ml right before measurement. Ideal measurement concentrations were determined by pre-testing the ideal particle per frame value (20-100 particles/frame). The following settings were chosen according to the manufacturer’s manual using the 488 nm laser with a camera level of 12, a slider shutter of 1,200, and a slider gain of 146. Per sample, five to six videos of each 60 s with 25 frames per s were captured at a constant temperature of 25 °C and assuming a water-like viscosity. Of the six captured videos, all videos passing the validity assessment by the software were analysed with a detection threshold of seven and blur size and maximum jump distance set to automatic. In addition, raw data was normalised against the leaf fresh weight of the individual experiment and further analysed using Microsoft Excel for Mac version 16.55 (21111400) by merging the raw data of separate biological replicates with the same treatment. Data were plotted in GraphPad (Boston, MA, USA) Prism version 8.

### Protein identification by mass spectrometry

Samples for mass spectrometry were prepared using the single-pot, solid-phase-enhanced sample-preparation (SP3) technology (Hughes *et al*. 2019) before being subjected to a trypsin digest over night at 37 °C. Samples were dimethyl labelled on peptide-level for 2h using 30 mM sodium cyanoborohydride and 30 mM ^12^CH_2_O for control and 30 mM ^13^CD_2_O for *B. hordei*-infected samples (Boersema *et al*. 2009). Reactions were quenched with 100 mM Tris after 2h labelling at 37°C, combined in a 1:1 ratio, and purified with custom-packed C18 StageTips (Rappsibler et al 2007). Peptides were analysed by nano HPLC-MS/MS with a Ultimate 3000 nanoRSLC (Thermo Fisher Scientific, Darmstadt, Germany) operated in a two-column setup (2 cm PepMap C18 trap, 75 µm ID, and 25 cm Acclaim PepMap C18 analytical column, 75 µm ID, Thermo Fisher Scientific, Darmstadt, Germany), coupled to a Bruker impactII Q-TOF instrument using gradient and acquisition parameters as described (Misas Villamil *et al*. 2019). Peptides were identified with the MaxQuant software package (Tyanova *et al*. 2016), version 1.6.10.43, using the *B. hordei* and *H. vulgare* UniProt reference proteomes (downloaded December 2019) including isoform-specific sequences and appended standard contaminants as database. For the query, up to one missed cleavage was allowed, and cysteine carboxyamidomethylation (57.0214 Da), lysine and N-terminal dimethylation (^12^CH_2_O 28.0313 Da; ^13^CD_2_O 34.0631 Da) were set as labels, and methionine oxidation as a variable modification. The false discovery rate for spectrum, peptide and protein identification was set to 0.01.

Only proteins observed in at both biological replicates and identified with at least two unique peptides were considered in further analysis. Transmembrane domains and signal peptides were predicted using TMHMM version 2.0 (Krogh *et al*. 2001); https://services.healthtech.dtu.dk/services/TMHMM-2.0/) and SignalP version 5.0 ((Almagro Armenteros *et al*. 2019); https://services.healthtech.dtu.dk/services/SignalP-5.0/), respectively.

### Protease protection assay

PME-derived EVs were resuspended in 20 mM Tris-HCl (pH 7.5). Proteinase K digest was performed as described previously (Chow *et al*. 2019). Samples were split into three equal parts and treated with proteinase K at a final concentration of 5 µg/ml, in the absence or presence of 0.05% Triton X-100. One part was kept untreated. All parts were incubated at 37 °C for 1 h prior to denaturing protein gel electrophoresis and immunoblot analysis. Alternatively, a protease protection assay (Rutter & Innes 2017) was performed. The EV sample was split into four equal parts and treated with 100 µg/ml trypsin, 5% Triton X-100 or pre-treated with 5% Triton X-100 followed by treatment with 100 µg/ml trypsin. Triton X-100 treatment was carried out on ice for 30 min. Proteins were digested with trypsin at 37 °C for 1 h. All samples, including the control sample, were subjected to the same incubation temperatures and times.

## Results

### The protein profile of barley leaf AWF changes in response to powdery mildew infection

To extract AWF, we established a procedure originally developed for barley (Rohringer *et al*. 1983) and further amended it according to a recently reported method ((Rutter & Innes 2017); Figure 1A – see Materials and Methods for further details). We collected AWF from non-inoculated (control) and *B. hordei*-inoculated primary barley leaves and visualised its protein content by sodium dodecyl sulfate polyacrylamide gel electrophoresis and silver staining. All AWF-derived protein samples (collected at 0, 24 and 72 hours post inoculation (hpi)) showed a markedly different protein pattern compared to a corresponding whole leaf extract sampled at 72 hpi (Figure 1B). Prominent protein bands of the whole leaf extract were mostly absent or underrepresented in the AWF samples; conversely, pronounced protein bands of the AWF samples were essentially unrecognizable in the whole leaf extract. We also noticed that the AWF protein profile changed in the course of *B. hordei* infection: Samples derived from AWF collected at 24 and 72 hpi showed overall additional and/or more intense bands as compared to the non-inoculated (0 hpi) AWF control sample (Figure 1B).

**Figure 1.**
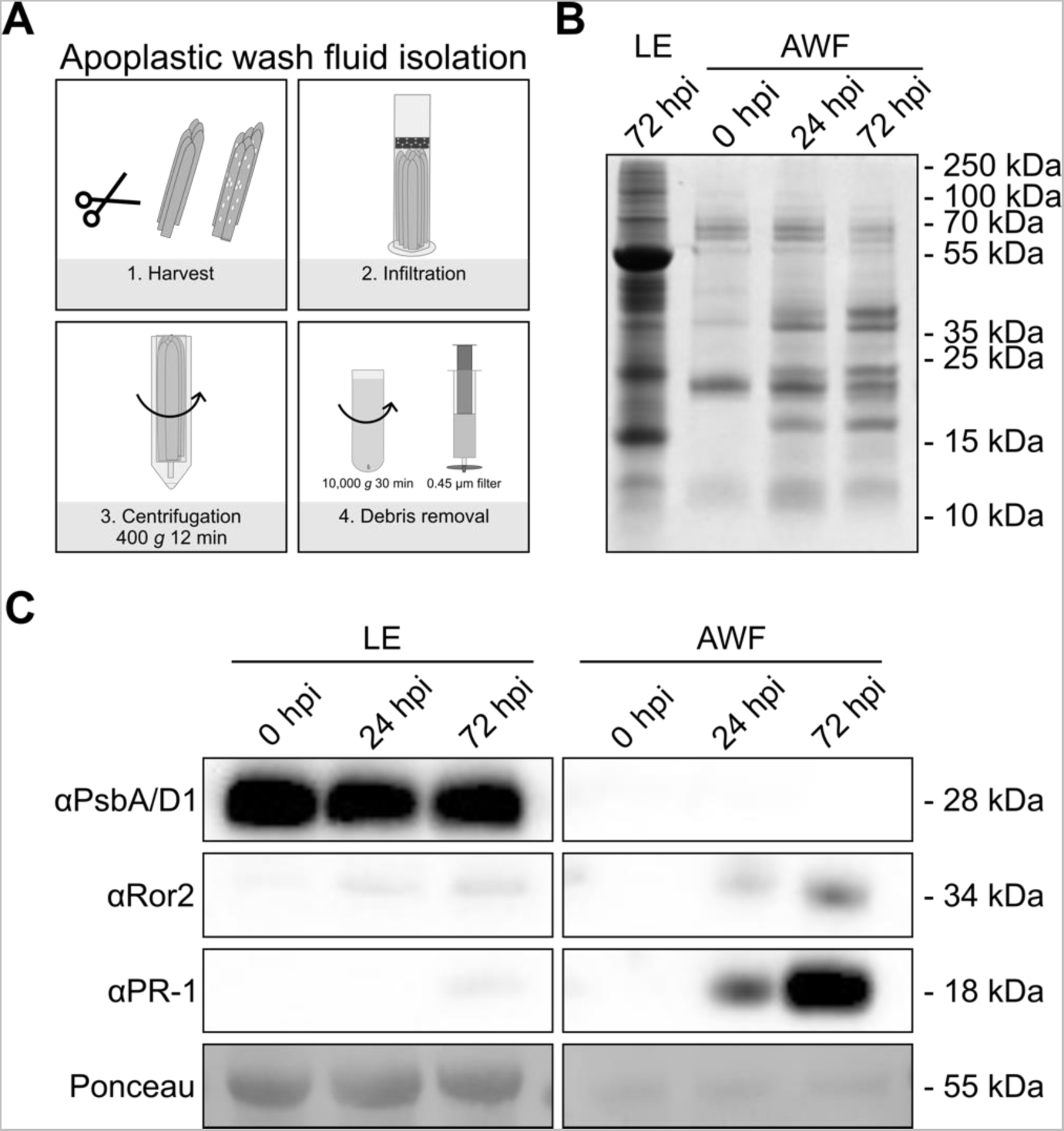
Challenge with *B. hordei* induces changes in the protein profile of the barley leaf AWF. **A** Scheme depicting the workflow for the extraction of AWF from barley primary leaves. The procedure is based on a previously described method (Rohringer *et al*. 1983) and involves leaf harvest, the infiltration of the leaf apoplastic space with buffer, collection of the AWF by centrifugation, and removal of the debris from the AWF by filtering and centrifugation. For further details, see Materials and Methods. **B** Leaf extract or AWF was collected from barley leaves that were sampled either prior to inoculation (0 hpi, 7-day-old) or at 24 hpi (8-day-old) or 72 hpi (10-day-old) with *B. hordei*. Proteins (0.25 µg per sample) were separated by gel electrophoresis and stained with silver. The experiment was performed in two independent biological replicates with similar outcome. **C** Immunoblot of total leaf extract (LE) and AWF probed with antibodies specific for the chloroplastic/cytosolic contamination marker PsbA/D1 (predicted molecular mass 28 kDa), the potential EV marker Ror2 (predicted molecular mass 34 kDa), and the secreted defence marker PR-1 (predicted molecular mass 18 kDa). Total leaf extract and AWF was collected from barley leaves that were sampled either prior to inoculation (0 hpi) or at 24 hpi or 72 hpi with *B. hordei*. Gels were loaded with 2.5 μg protein per sample. The apparent molecular masses of proteins (given on the right) were judged by comparison with protein standards analysed on the same gel. Staining with Ponceau S (the prominent band corresponding to the large subunit of RuBisCO) served to demonstrate equal loading. The experiment was performed in three independent biological replicates with similar results.

We further analysed samples of total leaf extract and AWF (collected at 0, 24 and 72 hpi) by immunoblot analysis. We probed the blots with antibodies directed against photosystem II protein D1 (PsbA/D1, a thylakoid membrane marker protein; (Huokko *et al*. 2021), Ror2 (a t-SNARE protein and the barley ortholog of the established Arabidopsis EV marker PEN1; (Collins *et al*. 2003)) and the defence-related protein PR-1 (pathogenesis-related protein 1; (Pečenková *et al*. 2022)). While we detected as expected strong bands using the αPsbA/D1 antibody in the leaf extract at all time-points indicated, the signal was below detection limit in all AWF samples, suggesting little, if any, chloroplastic/cytoplasmic contamination in the AWF (Figure 1C). This observation was supported by a comparison of unprocessed (non-infiltrated) leaves and leaves subjected to AWF isolation stained with trypan blue to visualise dead cells (Supplemental Figure1). The Ror2 protein was undetectable prior to pathogen challenge in both leaf and AWF samples; however, bands of weak (leaf extract) or medium (AWF) intensity were present in the samples collected at 24 and 72 hpi. In the case of the AWF-derived samples, signal intensity at 72 hpi was higher than at 24 hpi. A similar pattern was seen for the accumulation of PR-1: The protein was below detection limit in leaf extract collected at 0 h and 24 hpi and only weakly recognizable in the 72-hpi sample. In the AWF samples, the protein was not detectable at 0 hpi and increased in abundance from 24 hpi to 72 hpi, showing a strong signal at the latter time-point (Figure 1C). Based on the combined examination of protein extracts by protein gel electrophoresis (Figure 1B) and immunoblot analysis (Figure 1C) at two time points following inoculation with *B. hordei*, we conclude that (1) the modified extraction protocol yields AWF with minimised chloroplastic/cytosolic protein contamination, and that (2) the method is sensitive enough to detect pathogen-induced changes in the AWF-associated protein profile. Considering the pronounced changes in the AWF protein pattern seen at 72 hpi and accumulation of the PEN1 ortholog Ror2, we focused on this time-point for the following analyses.

### The barley leaf apoplast contains a heterogeneous population of prototypical EVs

To isolate crude EVs from barley leaf AWF, we compared two different methods. First, we used a sequential differential ultracentrifugation protocol to collect two EV fractions pelleted at 40,000 *g* (P40) and 100,000 *g* (P100; (Rutter & Innes 2017); Figure2A). Alternatively, we captured EVs by polymer-mediated enrichment (PME, (Grunt *et al*. 2020); Figure2B). To validate the presence of vesicle-like structures, we performed transmission electron microscopy on P40, P100, and PME fractions isolated from the AWF of non-inoculated and inoculated (sampled at 72 hpi) barley leaves. Electron microscopy samples were stained with uranyl acetate except the PME fraction, for which phosphotungstic acid was used, because uranyl acetate stained polymers used during isolation of EVs by PME (Supplemental Figure2).

The P40 fraction frequently contained particles of varying size with a characteristic cup shape (Figure2C). The cup shape is a known technical artefact of the negative staining of spherical lipid-bilayer compartments and is commonly used as an indicator for vesicles (Théry *et al*. 2006; Chernyshev *et al*. 2015). The presumed vesicles were accompanied by smaller granular particles that did not display any apparent lipid bilayer and smaller, less electron-dense particles (Figure 2C). Inoculation with *B. hordei* did not affect the morphology of the vesicles observed at 72 hpi, and smaller grainy objects were still sporadically present, while electron-light objects were largely absent (Figure2C). The small granular objects, which were less predominant in the P40 fraction, were the main constituent of the P100 fraction and characteristic cup-shaped vesicles were only occasionally visible (Figure2C). Circular objects present in the P100 fraction appeared larger than those collected from the P100 fraction of non-inoculated leaf tissue (Figure2C). Their granular appearance was reminiscent of ribulose-1,5-bisphophate carboxylase/oxygenase (RuBisCO) complexes previously documented in other species (Raunser *et al*. 2009; Bowien & Mayer 1978). However, we did not detect RuBisCO by immunoblotting of the P100 fraction (Supplemental Figure 3), essentially excluding that the circular objects represent RuBisCo complexes. Comparing the constituents recovered at 40,000 *g* and 100,000 *g*, we conclude that centrifugation at 40,000 *g* is sufficient to isolate vesicle-like structures from barley leaf AWF, while higher centrifugation force (100,000 *g*) yields mostly granular particles of unknown origin and identity. Particles isolated by PME mostly comprised characteristic cup-shaped vesicles of different size (Figure2D). Additionally, smaller electron-light particles similar to those observed in the P40 fraction and, rarely, small grainy particles were present. Infection by *B. hordei* (72 hpi), did not change the observed vesicle morphology compared to the P40 fraction (Figure2D). Together, these results demonstrate that PME-derived EVs largely resemble the centrifugation based P40 fraction. We, thus, conclude that characteristic vesicles that have cup-shaped morphology in electron microscopic analysis can be isolated from the barley leaf apoplast using two different methods.

**Figure 2.**
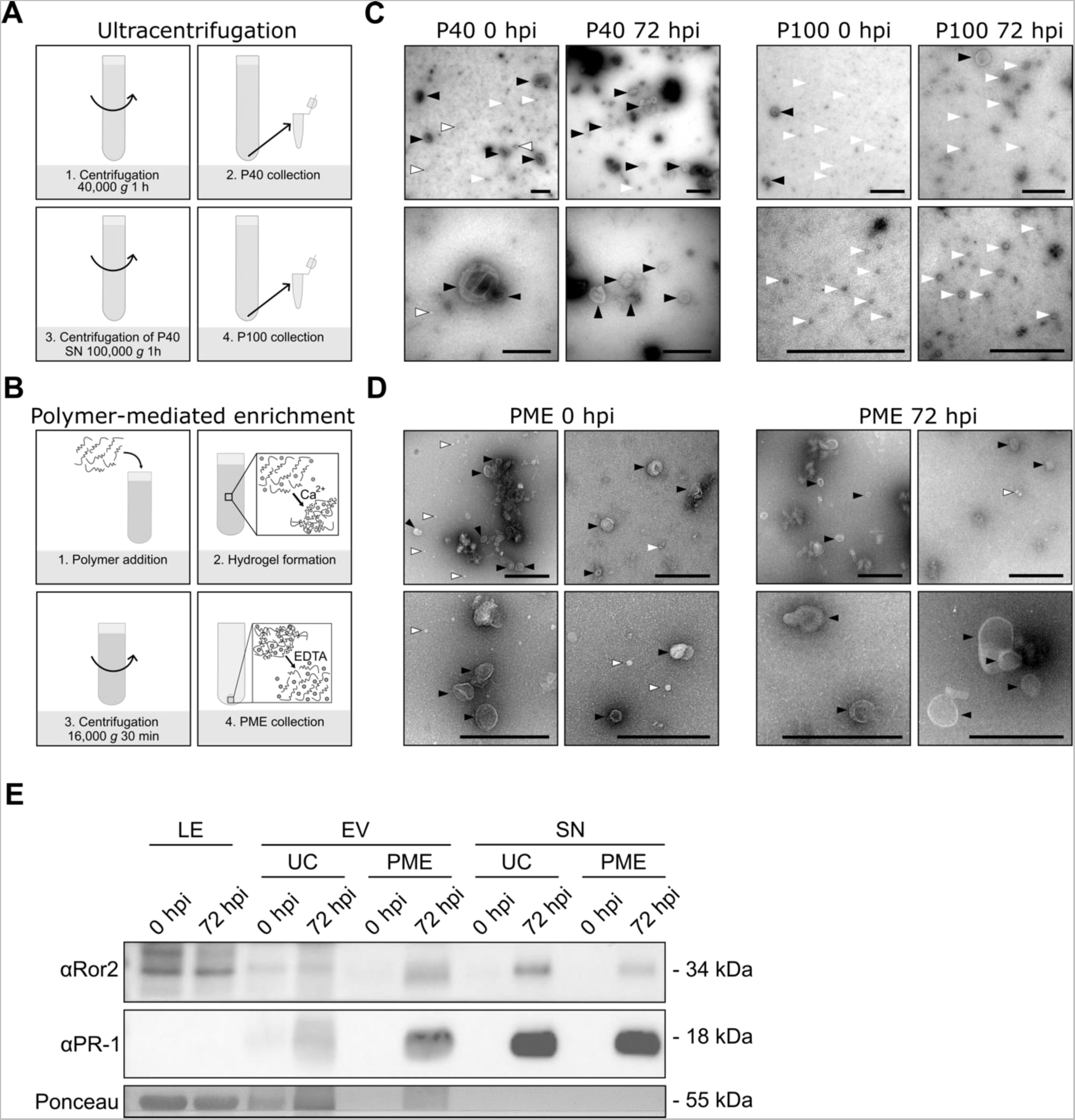
Characteristic cup-shaped structures are present in the P40 fraction and following PME of EVs isolated from the barley leaf AWF. **A** Scheme of the workflow for the isolation of crude EVs (P40 and P100) from barley AWF by differential ultracentrifugation at 40,000 *g* and 100,000 *g* (Rutter & Innes 2017). See Materials and Methods for further details. **B** Scheme of the workflow for the isolation of crude EVs from barley AWF by PME according to the manufacturer’s instructions. EVs are captured in a polymer-based hydrogel in the presence of calcium (Ca^2+^) ions and released by the addition of the Ca^2+^ chelator ethylenediaminetetraacetic acid (EDTA). See Materials and Methods for further details. **C** Transmission electron micrographs of P40 and P100 fractions isolated from AWF collected from barley leaves that were sampled either prior to inoculation (0 hpi) or at 72 hpi with *B. hordei*. EVs were stained using uranyl acetate. Representative images of three (non-inoculated) and four (inoculated) biological replicates are shown. Typical cup-shaped morphology – black arrowheads, small granular particles – white arrowheads, electron-light particles – white arrowheads with black outlines. Scale bars = 500 nm. **D** Transmission electron micrographs of EVs isolated *via* PME from barley leaves that were sampled either prior to inoculation (0 hpi) or at 72 hpi with *B. hordei*. EVs were stained using phosphotungstic acid. Representative images of four (non-inoculated) or three (inoculated) biological replicates are shown. Typical cup-shaped morphology – black arrowheads, small granular particles – white arrowheads, electron-light particles – white arrowheads with black outlines. Scale bars = 500 nm. **E** The barley ortholog of the Arabidopsis EV marker PEN1, Ror2, is present in EV and supernatant samples. Immunoblot of total leaf extract (LE), EV, and supernatant (SN) samples probed with αRor2 (potential EV marker, predicted molecular mass 34 kDa) and αPR-1 (secreted defence marker, predicted molecular mass 18 kDa). Leaf extract, EV and supernatant samples were isolated from leaves of 10-day-old barley plants that were either not inoculated (-) or sampled at 72 hpi with *B. hordei* (+). The gel was loaded with 5 μg protein of total leaf extract, completely resuspended ultracentrifugation (P40) pellets, 20 μl of PME pellets resuspended in 150 µl Tris buffer, and 10 μl supernatant sample. Staining with Ponceau S (the prominent band corresponding to the large subunit of RuBisCO) served to demonstrate equal loading of the gels. Apparent molecular masses of proteins (given on the right) were derived from a comparison with molecular mass standards analysed on the same gel. The experiment was repeated in two biological replicates with similar results.

We further compared EVs isolated from the same starting material by either ultracentrifugation or PME in immunoblots probed with αRor2, αPR-1 and αPsbA/D1. Overall, we observed similar protein patterns (Figure 2E), supporting the conclusion that comparable EV populations can be isolated with both methods. At 72 hpi, Ror2 was detectable in both P40- and PME-derived EV preparations, with a stronger signal in the PME than in the P40-based sample. Interestingly, Ror2 was also detectable in supernatant samples (the EV-depleted AWF). There, the opposite trend was seen (Figure 2E). This indicates that extracellular Ror2 is both associated with EVs and present in the vesicle-free fraction. Surprisingly, PR-1 was also detected in both EV preparations following pathogen challenge, with a stronger signal in the PME-derived EV sample. Moreover, we observed a stronger signal for PR-1 in the supernatant of both centrifugation- and PME-derived EVs, indicating that the majority of this protein is secreted *via* the canonical pathway, i.e. not associated with EVs. The band corresponding to RuBisCO, visible after Ponceau staining of the proteins blotted on the membrane, suggests that there is some intracellular contamination in both displayed EV preparations; however, it is weaker in the PME sample. Together with the electron micrographs (Figure 2C and D), the presented data indicate that PME is better suited than ultracentrifugation to enrich EVs and their associated proteins (e.g. Ror2), and to reduce intracellular contaminants related to the isolation procedure.

Next, we determined the size (hydrodynamic diameter) and abundance of particles recovered in ultracentrifugation- and PME-based EV preparations *via* NTA. NTA profiles obtained for P40-, P100- and PME-derived EVs each revealed a typical size distribution pattern for polydisperse samples (Vogel *et al*. 2021), i.e. a broad distribution with several peaks (Figure 3). A main peak between 100 nm and 300 nm was evident in the P40 and PME samples and became more prominent in samples from infected leaves, which was also reflected by a strong increase in frequency of these in PME samples (Figure 3A, C, and D). In addition, a second, more moderate peak of particles between 400 nm and 500 nm in size became more prominent in PME samples following challenge with *B. hordei*, which might represent a second pathogen-responsive EV population (Figure 3C, Supplemental Figure 5B). Interestingly, the size distribution of P100 samples derived from non-inoculated tissue largely resembled the distribution of the corresponding P40 preparation, likely because the small circular objects observed in the P100 sample in electron microscopy (Figure 2C) were below the operation range of the NTA device (Figure 3B). Alternatively, the results might indicate that the centrifugation time at 40,000 *g* is insufficient to pellet dense particles. However, we found that longer centrifugation time rather affects particle integrity instead of improving their separation, in line with observations from other systems (Supplemental Figure 4; (Taylor & Shah 2015)). We noticed that in PME samples obtained from both non-inoculated and inoculated leaves, the harvest of particles per ml/g fresh weight was higher compared to the fraction isolated by centrifugation, and the associated variance was also lower (compare Figure 3A and 3B with Figure 3C). This outcome further substantiates that PME is more efficient and reproducible than high-speed centrifugation to enrich EVs from barley leaf AWF. In summary, we conclude that the barley leaf apoplast contains a heterogenous population of EVs with a main population of ∼100 nm - 300 nm in diameter. Further, this population rises after infection with *B. hordei*, suggesting a selective response of barley to the fungal pathogen.

**Figure 3.**
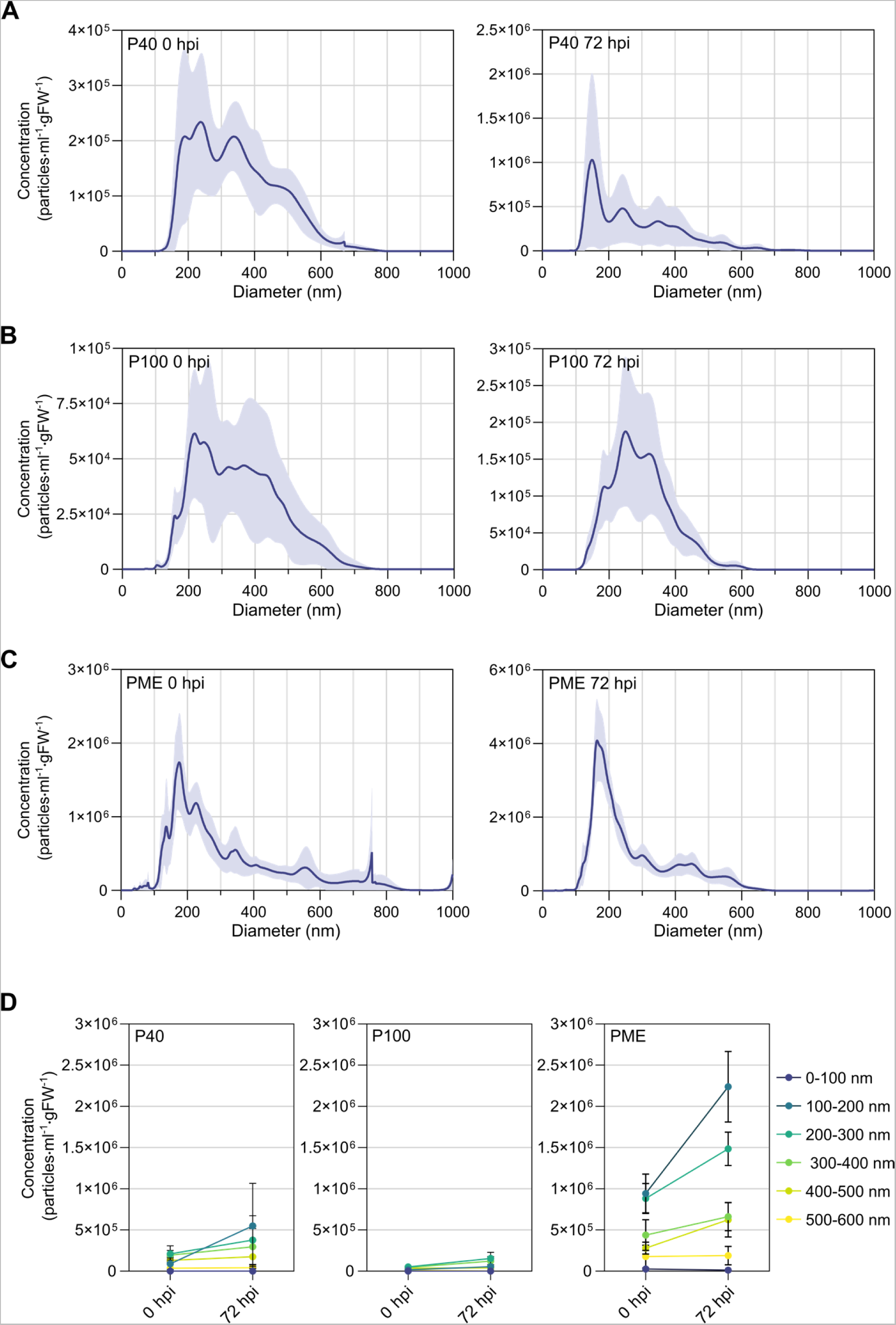
Barley leaf apoplastic EVs constitute polydisperse populations that are selectively responsive to infection with *B. hordei*. **A, B, C** NTA data showing the size distribution and mean concentration within the upper and lower 95 % confidence intervals of P40- (**A**), P100- (**B**), and PME-derived (**C**) EV samples. EVs were isolated from leaves of 10-day-old barley plants that were either sampled prior to inoculation (0 hpi) or at 72 hpi with *B. hordei*. Plots are based on data from five (P40, P100) or six (PME) independent biological replicates. **D** Comparison of EV concentrations in P40, P100 and PME samples from non-inoculated (0 hpi) and *B. hordei*-infected (72 hpi) plants, divided into 100 nm bins as indicated by the colour-coded legend on the right side of the plots. Points represent the mean, and error bars the standard deviation. Plots are based on data from five (P40- and P100-derived EVs) or six (PME-derived EVs) independent biological replicates. The plots are based on the dataset shown in panels **A**, **B** and **C** above.

### Evidence for the dynamic accumulation of separate barley leaf EV populations during fungal pathogenesis

Next, we investigated how barley leaf EVs accumulate throughout the full course of infection by isolating EVs *via* PME either prior to inoculation (0 hpi) or at 8, 24, 48, 72, 96, and 120 hpi with *B. hordei*. Their overall abundance increased as fungal proliferation progressed, in particular at 96 hpi and 120 hpi, reaching an overall abundance of almost 25-fold at 120 hpi in comparison to vesicles from non-inoculated control samples (Figure 4A, Supplemental Figure 5A). While the size distribution profile remained largely consistent over the time-course of infection, a major peak became apparent as the infection progressed (Figure 4A, B; Supplemental Figure 5A). Dividing the data into 100 nm bins revealed that this peak is caused primarily by a notable increase in particles ranging from 100 nm to 200 nm and 200 nm to 300 nm in diameter (Figure 4B). Vesicles of these two size classes did not change drastically until after 72 hpi (∼1- to 2.5-fold), but rose dramatically ∼10- and 50-fold 96 hpi and 120 hpi, respectively, compared to non-infected controls. Vesicles ranging from 300 nm to 400 nm and 400 nm to 500 nm in diameter showed a similar dynamic , but accrued much lower levels at 96 hpi and 120 hpi (∼2- to 3-fold compared to the non-infected control sample). The differential accumulation profile suggests that they might represent a separate EV population compared to the main peak caused by vesicles of 100 nm to 300 nm in diameter (Supplemental Figure 5B).

**Figure 4.**
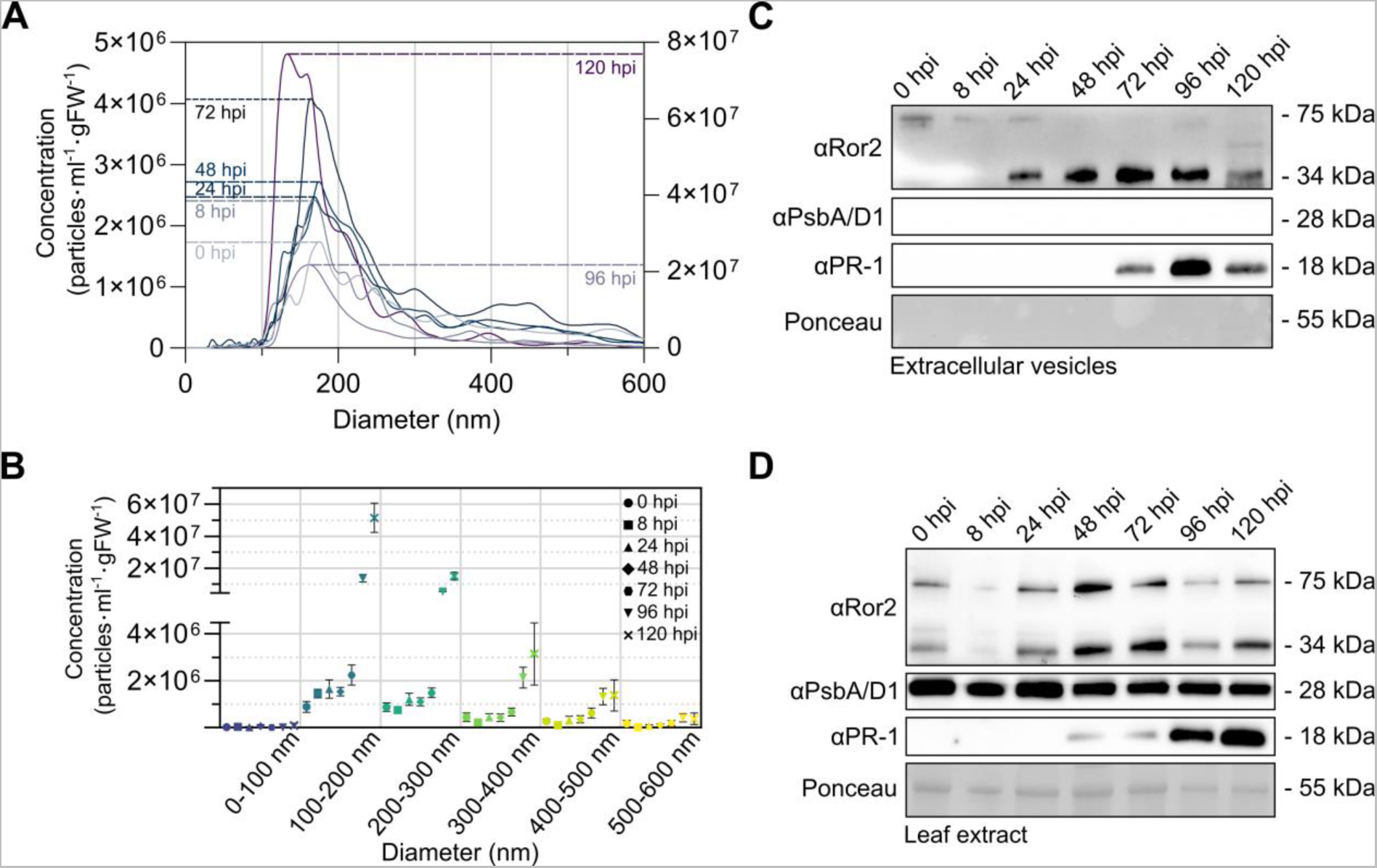
Levels of barley leaf apoplastic EVs and its associated candidate marker protein Ror2 increase highly during fungal infection. **A** NTA data showing the size distribution and mean concentration of PME-derived EVs isolated from barley leaves either prior to inoculation (0 hpi) or at 8, 24, 48, 72, 96 or 120 hpi with *B. hordei*. Note the different scales used: data for time points 0, 8, 24, 48, and 72 hpi refer to the left y-axis, data for 96 hpi and 120 hpi refer to the right y-axis. **B** Concentration of EVs displayed by 100 nm size fractions sampled at different time points after inoculation with *B. hordei*. Points represent the mean, error bars 95% confidence intervals. The experiment was performed in six independent biological replicates for time points 0-72 hpi and three independent biological replicates for 96-120 hpi. The graph is based on the dataset shown in A. **C, D** Immunoblots probed with antibodies specific for the potential EV marker protein Ror2 (predicted molecular mass 34 kDa), the secreted defence marker PR-1 (predicted molecular mass 18 kDa), and the chloroplastic/cytosolic marker PsbA (predicted molecular mass 28 kDa) in PME-derived EVs (**C**) or total leaf extract (**D**) at different time points after inoculation with *B. hordei*. Gels were loaded with 20 µl of EV pellets resuspended in 150 µl Tris buffer (pH 7.5) and mixed with 6x loading dye. For total leaf extract, 5 µg protein were loaded on the gel. Molecular masses of proteins (given on the right) were estimated from comparison to a molecular mass standard analysed on the same gel. Proteins on the blots were stained with Ponceau S (the prominent band corresponding to the large subunit of RuBisCO) and this served to demonstrate equal loading. The experiment was performed in six independent biological replicates for time points 0-72 hpi and three independent replicates for time points 96-120 hpi.

### Ror2 partly associates with EVs

To complement the NTA data, we monitored the accumulation of the potential EV marker protein Ror2 in vesicle, supernatant and total leaf extract samples by immunoblot analysis in the course of fungal infection. In the vesicle samples, monomeric Ror2 (predicted molecular mass of 34 kDa) was below detection limit prior to pathogen challenge (0 hpi), but from 24 hpi onwards, its levels increased steadily as fungal infection progressed (Figure 4C). Besides the monomeric Ror2 species, an additional specific Ror2 signal at a molecular mass of around 75 kDa was detectable in vesicle samples derived from non-inoculated leaves (0 hpi). This signal gradually decreased in intensity, reaching detection limit at later stages of fungal infection (from 48 hpi onwards). The 75-kDa Ror2 variant could also be recognized in total leaf extract of barley wild type, but not of *ror*2-1 mutant plants (Supplemental Figure 5C). In total leaf extract, both Ror2 signals (34 kDa and 75 kDa) remained essentially unaltered in intensity in the course of fungal infection (Figure 5D). In the vesicle samples, the 75 kDa species was resistant to treatment with up to 4M urea (Supplemental Figure 5D). In contrast to the vesicle samples, this variant was below detection limit in the corresponding supernatant samples, which nonetheless revealed a similar dynamics of the monomeric (34 kDa) Ror2 signal as seen in the respective vesicle samples in the course of fungal infection (Supplemental Figure 5E). Taken together, the data suggests that Ror2 in part associates with EVs. It further hints at opposing accumlation dynamics of monomeric Ror2 and a 75-kDa Ror2 variant in the vesicles upon challenge with *B. hordei*.

**Figure 5.**
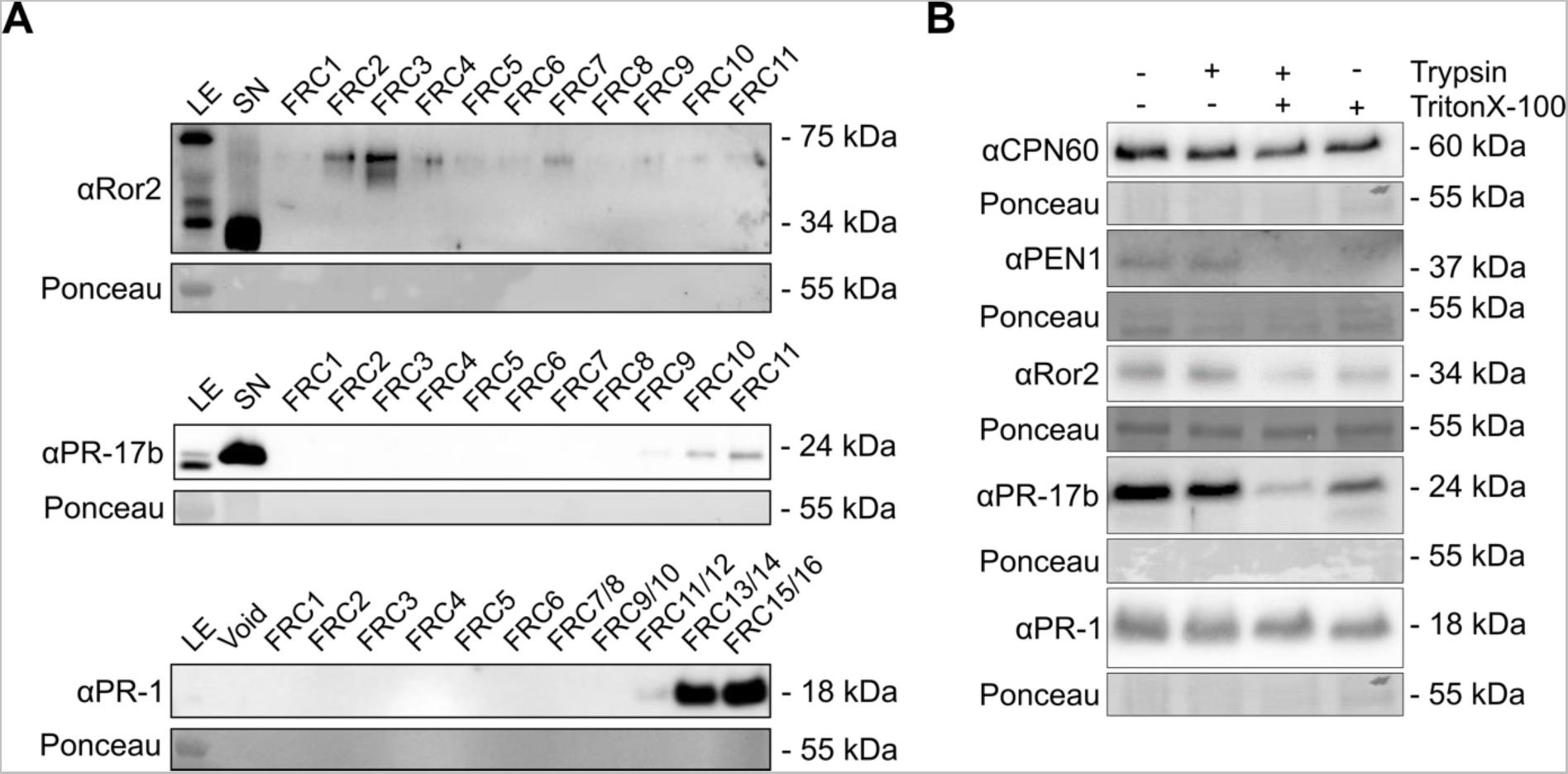
Ror2 might qualify as an EV marker. **A** Immunoblots probed with antibodies specific for the potential EV marker protein Ror2 (predicted molecular mass 34 kDa), the defence protein PR-17b (predicted molecular mass 24 kDa), and the secreted defence marker PR-1 (predicted molecular mass 18 kDa) in total leaf extract (LE), supernatant (SN) or void, and 11-16 fractions collected by size exclusion chromatography (SEC) of PME-derived EVs derived from barley leaves at 72 hpi with *B. hordei*. Gels were loaded with 10 µl supernatant, and/or 166 µl of each SEC fraction (FRC) or SEC void volume (after lyophilisation and resuspension in 10 µl) and mixed with 6x loading dye. For LE, 5 µg protein were loaded on the gel. Molecular masses of proteins (given on the right) were judged according to a molecular mass standard run on the same gel. Staining with Ponceau S (the prominent band corresponding to the large subunit of RuBisCO) served to demonstrate successful transfer. The experiment was performed in three independent biological replicates for each protein. **B** Immunoblots probed with antibodies specific for the potential EV marker protein CPN60 (predicted molecular mass 60 kDa), the potential EV marker protein Ror2 (predicted molecular mass 34 kDa), the Arabidopsis EV marker PEN1 (predicted molecular mass 37 kDa), the defence protein PR-17b (predicted molecular mass 24 kDa), and the secreted defence marker PR-1 (predicted molecular mass 18 kDa) in PME-derived EVs in barley leaves sampled at 72 hpi with *B. hordei*. EVs were treated with either trypsin or Triton X-100 or pre- treated with TritonX-100 followed by trypsin. Gels were loaded with 20 µl of treated EV pellets and mixed with 6x loading dye. Molecular masses of proteins (given on the right) were judged according to a molecular mass standard run on the same gel. The experiment was performed in seven independent biological replicates for Ror2 and PEN1, and three independent biological replicates for CPN60, PR-17b, and PR-1.

### The barley leaf apoplastic EV proteome contains many stress-related proteins and is only in part responsive to *B. hordei* infection

To better understand the proteins associated with EVs derived from the barley leaf apoplast and to explore potential EV markers, we isolated crude EVs from AWF from non-inoculated barley leaves and leaves sampled at 72 hpi by ultracentrifugation and examined them using label-free mass spectrometry. Our analysis identified in total 216 proteins based on two replicates, consisting each of a non-inoculated and an inoculated sample. Seventy of these proteins were present in both replicates and had a unique peptide count of at least two (Table 1). Interestingly, two of the three most abundant proteins were RuBisCO large subunit-binding proteins (subunits alpha and beta; A0A8I6W9H5 and A0A8I6YA55, respectively). RuBisCO large subunit-binding proteins are proteins of the chaperonin 60 family. The detection of cytosolic proteins is not unexpected as molecules associated with EVs to some extent represent the cytosolic content of their donor cells (Hurwitz *et al*. 2016). Moreover, the human chaperonin 60 (CPN60/HSP60) is a marker of exosomes, particularly in cancer cells (Caruso Bavisotto *et al*. 2017). The barley genome encodes two CPN60 isoforms that share high amino acid sequence similarities with human CPN60 (Supplemental Figure 6A), which is also reflected by comparable predicted 3D structures of the Arabidopsis, barley and human proteins (Supplemental Figure 6B). Using an antibody directed against Arabidopsis CPN60 subunit alpha we confirmed the accumulation of barley CPN60 in EVs in comparison to AWF or supernatant, but not compared to total leaf extract (Supplemental Figure 6C). Further marker candidates are the two 14-3-3 domain-containing proteins (M0X3R2, F2CRF1), given that 14-3-3 proteins have been proposed to represent markers of human exosomes (Choi *et al*. 2015).

**Table 1.**
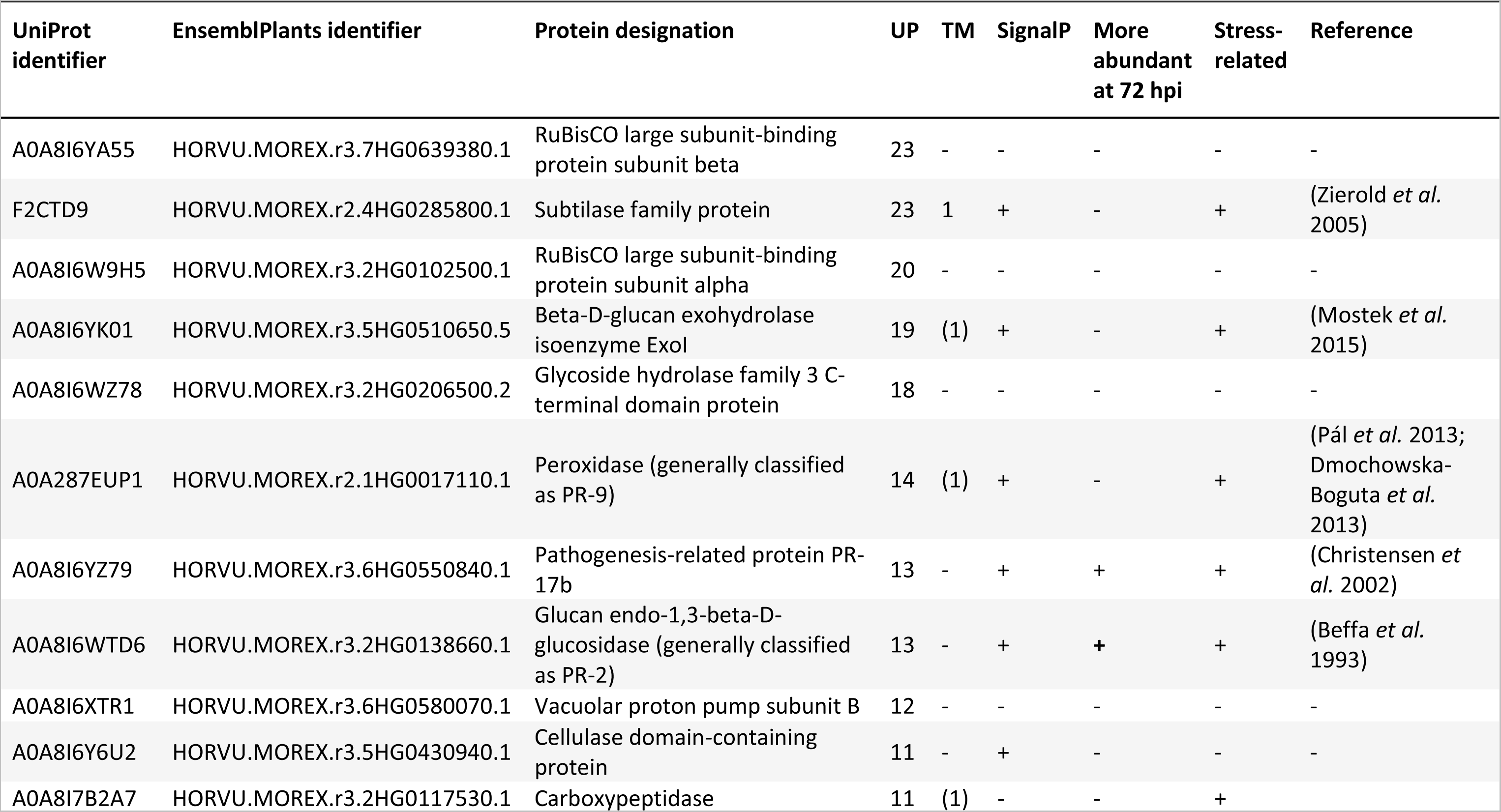
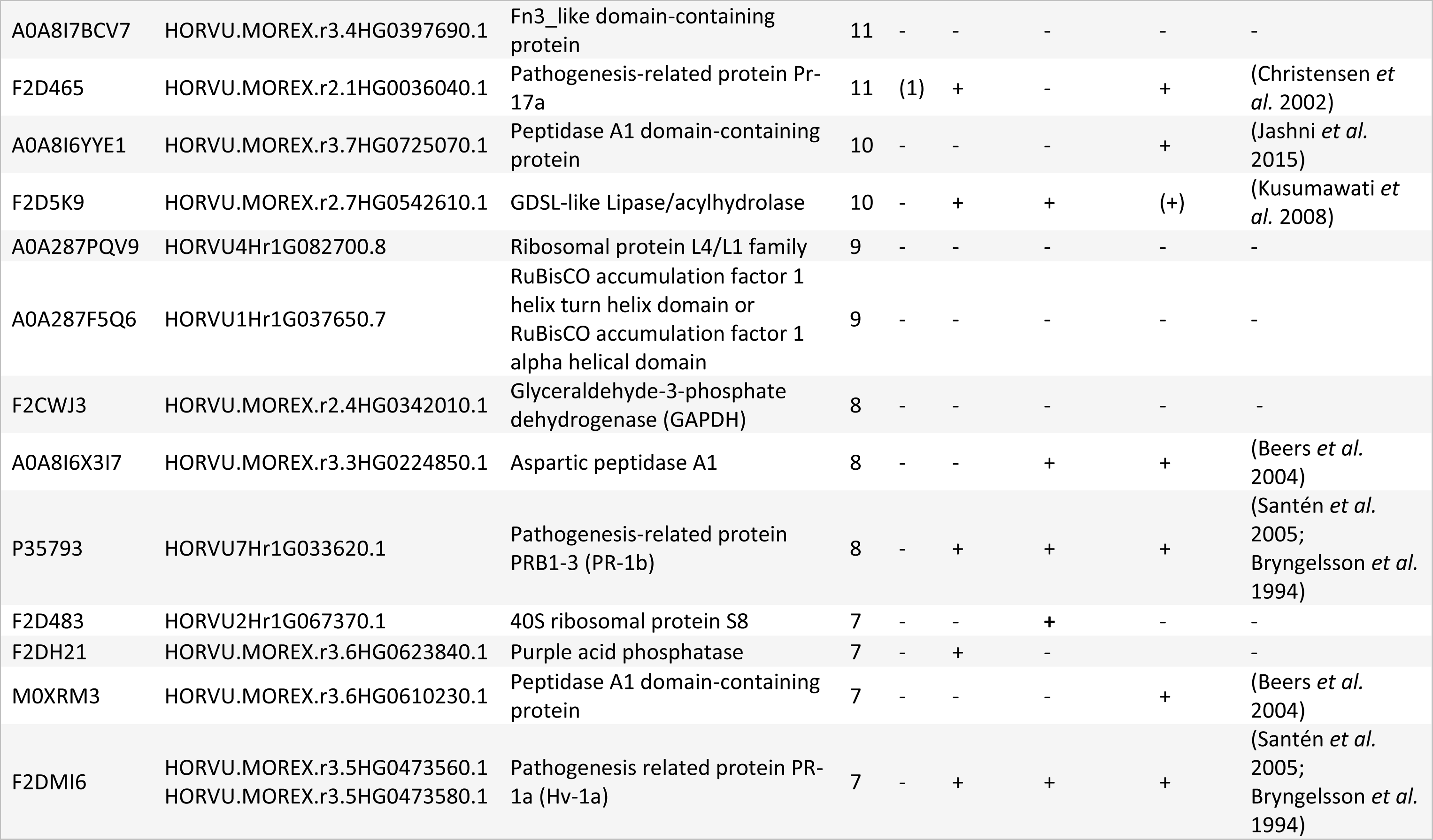
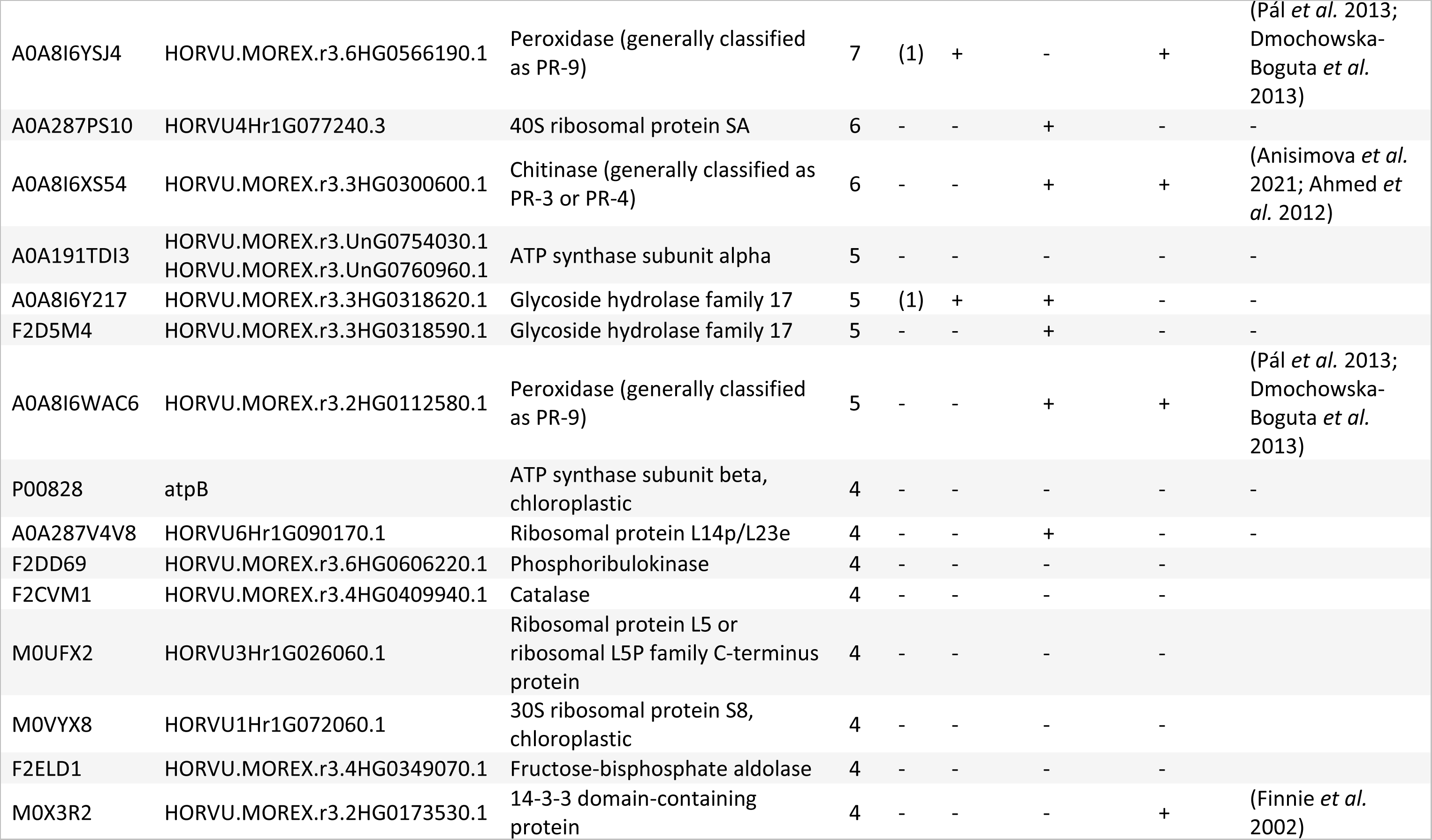
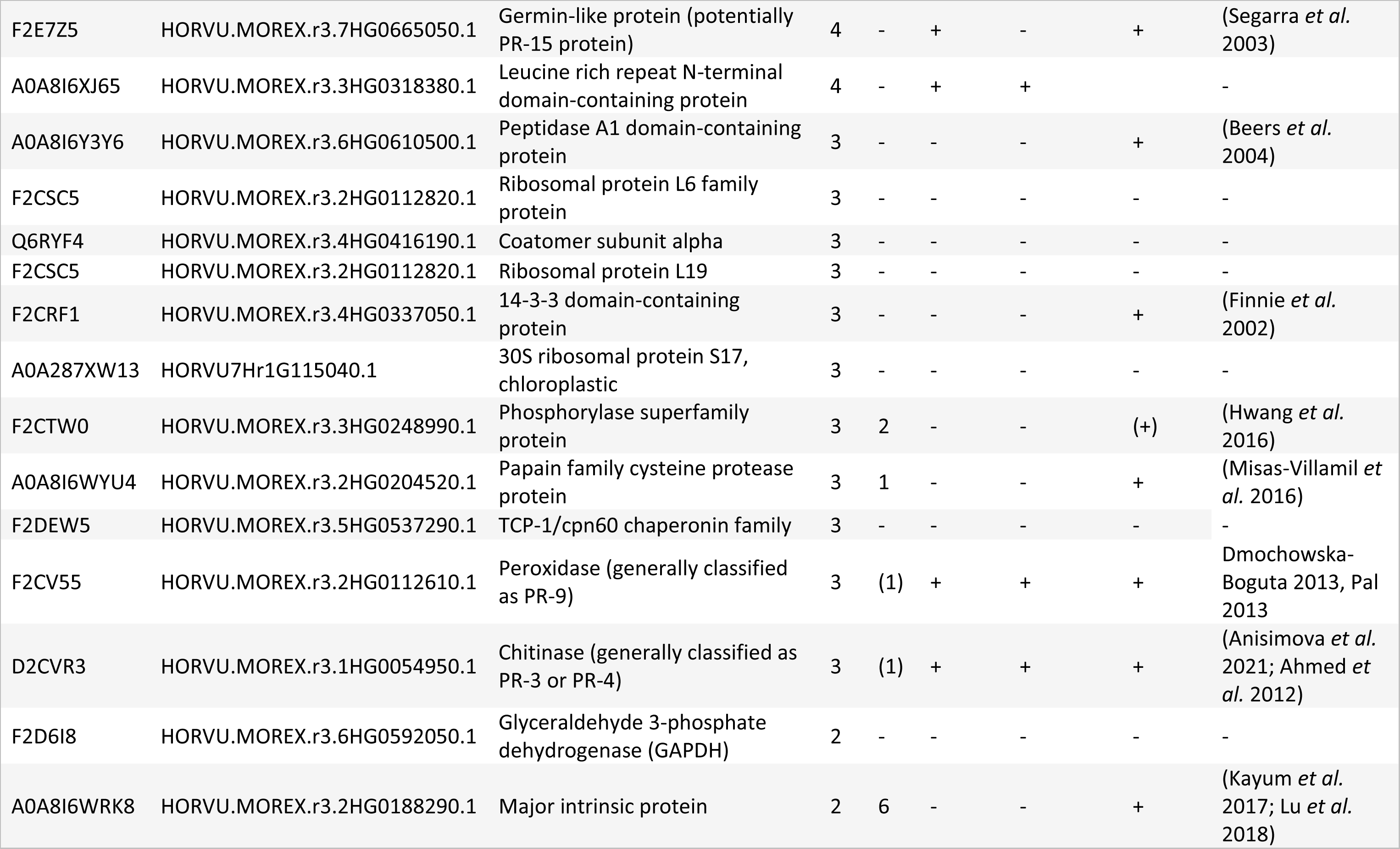
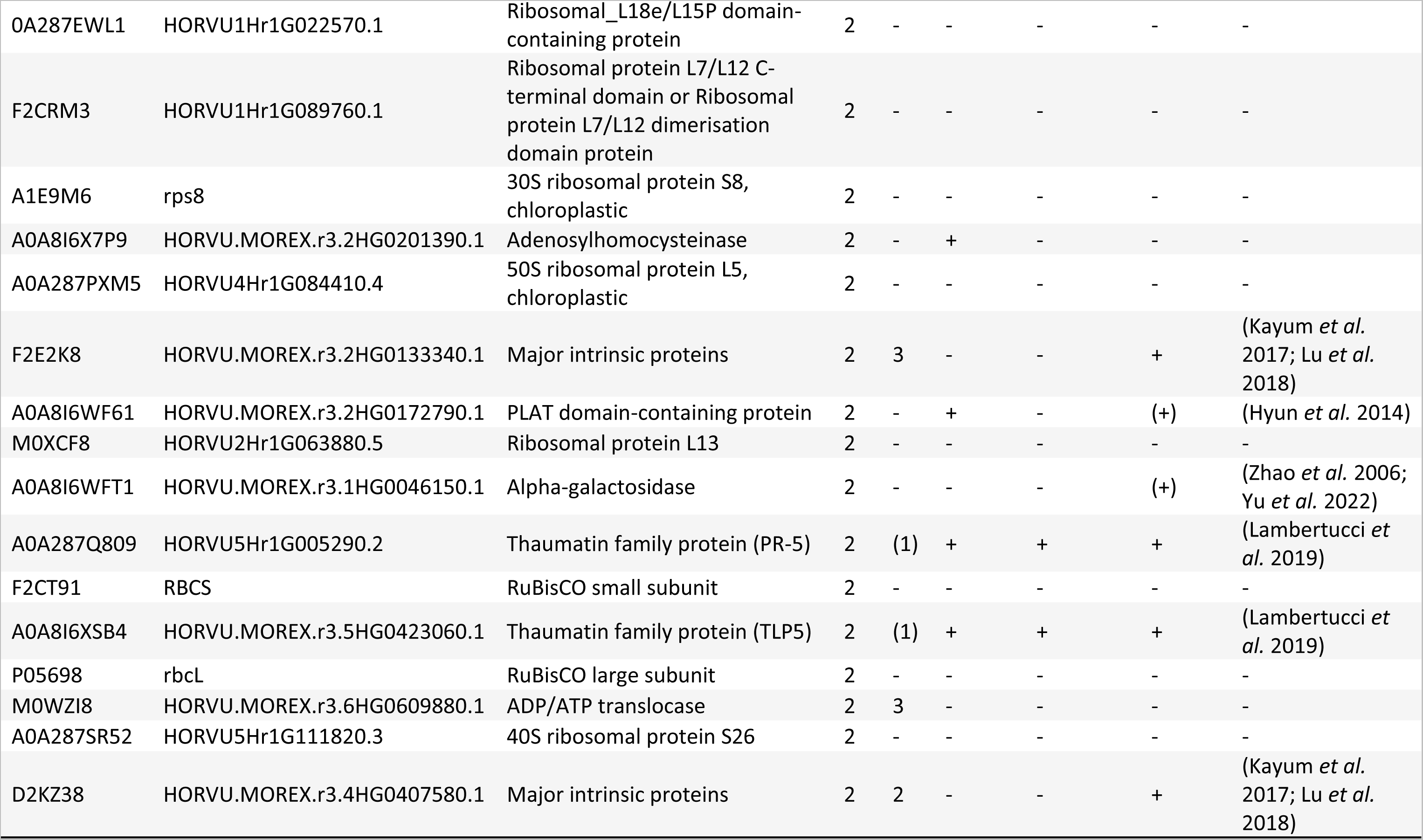
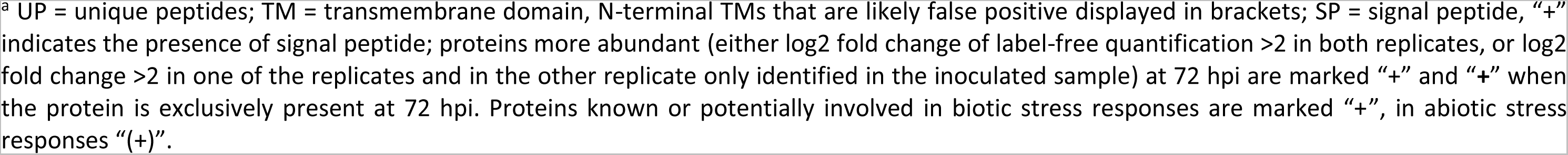
Proteins identified by mass spectrometry that were represented by at least two unique peptides and detected in both biological replicates ^a^.

Other noteworthy proteins detected in the crude EV proteome are cell wall remodelling enzymes (A0A8I6YK01, A0A8I6WZ78, A0A8I6Y6U2, A0A8I6Y217, and F2D5M4) that might aid the reorganisation of the extracellular space for the passage of EVs. In line with previous publications investigating plant EV proteomes (Rutter & Innes 2017; Regente *et al*. 2017), we found 31 proteins (∼22% of all identified proteins) that are known to or likely to be involved in either biotic or abiotic stress responses (Table 1), some of which have already been described in the barley-*B. hordei* interaction (e.g. PR-5 (A0A287Q809, (Lambertucci *et al*. 2019)). It is striking that many of the stress-related proteins can be classified as pathogenesis-related proteins (13 proteins, ∼19% of all identified proteins, Table 1). This is surprising since most pathogenesis-related proteins have an amino-terminal signal peptide and are delivered to the extracellular space *via* the conventional secretion pathway. However, the total number of proteins with a predicted signal peptide (∼30% of all identified proteins, Table 1) was moderate. Further, we identified several proteins that are reported to be secreted but lack a canonical amino-terminal signal peptide such as a carboxypeptidase (A0A8I7B2A7) and A1 peptidases (A0A8I6YYE1, A0A8I6X3I7, M0XRM3, and A0A8I6Y3Y6; (Kusumawati *et al*. 2008; Agrawal *et al*. 2010; Segarra *et al*. 2003)). In summary, these results suggest EVs may be mediators of a distinctive secretion pathway in barley, especially for stress-related proteins. Alternatively, these proteins may loosely associate and thus co-purify with EVs.

In the samples collected at 72 hpi, 18 proteins had a log2 fold change >2 in the label-free quantification compared to samples derived from uninfected tissue, or were only detected in samples from infected tissue (Table 1). Among these, the portion of stress-related proteins was even larger compared to non-inoculated tissue (12 proteins, ∼66%), in particular the fraction of already characterised or probable pathogenesis-related proteins (10 proteins, ∼55%). We found PR-1a and PR-1b, also associated with P40 and PME EV preparations as well as the supernatant (Figure 2E, Supplemental Figure 6D); interestingly PR-1a and PR-1b often serve as markers for conventionally secreted proteins in immunoblots (Wang & Fobert 2013). Only two proteins were uniquely present after inoculation: the 40S ribosomal protein S8 (F2D483) and a β-1,3-glucanase (A0A8I6WTD6), which is classified as member of the PR-2 proteins in the context of plant-microbe interactions (Dos Santos & Franco 2023). Together with the NTA results (Figure 3 and 4), our findings indicate that the *B. hordei*-induced changes are rather quantitative than qualitative, which is in line with a previous study investigating Arabidopsis EVs upon infection with the bacterial pathogen *Pseudomonas syringae* (Rutter & Innes 2017).

### Ror2 might qualify as an EV marker

Finally, we wanted to assess whether proteins identified by mass spectrometry in crude P40 samples as well as the barley orthologue of Arabidopsis EV marker PEN1, Ror2, are directly associated with barley leaf EVs. We probed size exclusion chromatography fractions and protease-treated EV samples using the antibody raised against Ror2 (Supplemental Figure 5E) and antibodies directed against PR-1 and PR-17b. The latter (A0A8I6YZ79) was represented by 13 unique peptides and hence one of the most abundant proteins isolated from the AWF of inoculated leaves. Additionally, we selected PR-1, as surprisingly both PR-1a (F2DMI6) and PR-1b (P35793) were detected by protein mass spectrometry. Size exclusion chromatography is a common strategy to purify crude EV samples based on separation from soluble proteins and other small molecules by size. Here, we used this chromatographic technique to provide evidence of whether a protein is firmly associated or just co-isolated with EVs. We observed that EVs primarily elute in chromatography fractions 2-4, and the NTA profile of the fraction with the highest concentrations of EVs mirrored that of crude EVs (Supplemental Figure 7). Only the ∼75-kDa Ror2 signal described above (cf. Figure 4C) was observed in the fractions even though the monomeric signal (34 kDa) is present in the supernatant of the same sample (Figure 5A). This signal was most abundant in fractions 2-4, which coincide with EV elution. Additionally, it was detected in fraction 7, suggesting the presence of different pools of Ror2 present in crude EV samples. Both PR-1 and PR-17b eluted in later fractions (PR-1 in fractions 11-16, PR-17b in fractions 10-11) where soluble proteins are expected, indicating that these proteins are co-isolated, likely due to aggregate formation or technical artefacts, possibly arising from their high abundance in the apoplast, or a loose attachment to EVs (Figure 5A).

Protease protection assays are valuable tools to determine if a protein resides in the EV lumen or outside (Chow *et al*. 2019; Chaya *et al*. 2023; Théry *et al*. 2018). Equal amounts of PME-derived EV samples were treated with either proteinase K or trypsin in the presence or absence of Triton X-100, or were left untreated. Proteins within the EV lumen are expected to be protected by the lipid membrane and remain intact after exposure to the protease but to be digested when detergent (Triton X-100) is added. Surprisingly, the results differed between the protocols applied (trypsin or proteinase K). Ror2 was partially digested only when both trypsin and the detergent were present, suggesting that Ror2 is present in the EV lumen (Figure 5C). This notion is supported by the observation that the Ror2-specific ∼75- kDa signal elutes in SEC fractions together with EVs (Figure 5A). On the other hand, when proteinase K was applied, Ror2 could no longer be detected once proteinase K was included (irrespective of the presence or absence of Triton X-100), indicating that the t-SNARE protein is not protected from digestion and, thus, is unlikely to reside in the EV lumen (Figure 5B). In addition to the Ror2-specific antibody, we also used an antibody directed against PEN1, the ortholog of Ror2 in *A. thaliana*. This antibody cross-reacts with the PEN1 ortholog of the grass species *Sorghum bicolor* (Chaya *et al*. 2023). We also observed cross-reaction of the αPEN1 antibody with barley Ror2. In the protection assay, the signal generated by the αPEN1 antibody was protected from digestion with trypsin in the absence of Triton X-100 (Figure 5C). However, in three out of four replicates, we detected no Ror2 signal in the detergent-only control when αPEN1 was used, hinting at a possible interference of Triton X-100 with this antibody and/or the activation of a vesicle-resident protease in the presence of the detergent. Similar to Ror2, in the absence of the detergent PR-17b was also protected in the protection assay with trypsin but digested in the assay with proteinase K (Figure 5C). Together with the SEC results described above (Figure 5A), the former hints at a potentially loose connection of PR-17b to EVs. As for the αPEN1 antibody, though less pronounced, we noticed a reduction in signal intensity for PR-17b in the sample with the detergent but lacking trypsin (Figure 5C). Interestingly, PR-1 appeared to be completely resistant to both trypsin and proteinase K digestion, as a signal with unaltered intensity could still be detected when Triton X-100 was added, even though both barley PR-1 proteins (PR-1a and PR-1b) possess multiple cleavage sites for both proteases (Supplemental Figure 8). CPN60 was only tested with trypsin but also seemed to be protected, making it an additional EV marker candidate next to Ror2.

## Discussion

Previous studies have highlighted the involvement of EVs in interactions with various microbes, ranging from pathogenic fungi to mutualistic bacteria (Cai *et al*. 2018; Li *et al*. 2022). Here, we present evidence demonstrating the pathogen-induced accumulation of EVs in the interaction with an obligate biotrophic fungal pathogen. Utilizing transmission electron microscopy, NTA and proteomic analyses, we conducted an in-depth characterisation of EVs obtained from the apoplast of both non-inoculated and inoculated barley leaves using two distinct isolation methods. Currently, differential ultracentrifugation is the primary method of choice for plant EV isolation, although challenging to establish in systems where upscaling is difficult (Rutter & Innes 2017; Cai *et al*. 2018; Regente *et al*. 2009). The PME procedure and differential ultracentrifugation revealed EVs with similar quality and morphological characteristics (Figure 2, Figure 3), suggesting that both methods are equally suited for EV isolation. This facilitated a time-course experiment, which revealed that barley responds to infection by the powdery mildew fungus with an increase in apoplastic EV abundance. These pathogen-responsive EVs likely comprise at least two distinct EV populations that vary in size and accumulation pattern (Figure 4, Supplemental Figure 5). As recently reported for EVs isolated from the monocot sorghum, the barley EV proteome was found to be enriched in stress-related proteins, especially following inoculation (Table 1). This finding, coupled with the aforementioned evidence for population-specific accumulation profiles (Figure 4, Supplemental Figure 5), suggests a biological function of EVs in the interaction of barley with *B. hordei*.

The apoplastic space is a well-known battleground in plant-microbe interactions (Du *et al*. 2016; Doehlemann & Hemetsberger 2013), including the challenge of barley by powdery mildew (Felle *et al*. 2004; Sargent 1977). The vesicles in this location may play roles in the outcome of the interaction. A key objective of this study was thus to establish a reliable and reproducible isolation procedure for barley leaf AWF-derived EVs. For this, to observe inoculation-induced changes in the apoplast, leaf-derived AWF with minimal cytosolic contamination is essential as starting material. Matching this criterion is pivotal for an accurate study of the vesicles, as liposomes resulting from cellular membrane damage can confound genuine EV analysis (Théry *et al*. 2018). We adapted existing protocols (Rutter *et al*. 2017; Rohringer *et al*. 1983) to achieve a reduction of cytosolic contamination below detection limits (Figure 1). Moreover, we confirmed differential protein accumulation, e.g. of the canonical stress response protein PR-1, in the barley apoplast upon inoculation with the powdery mildew pathogen, emphasizing the presumed significance of the extracellular defence response after challenge with *B. hordei* (Figure 1; (Tamás *et al*. 1997)). Employing the established protocol based on ultracentrifugation (Rutter & Innes 2017) alongside a novel polymer-based approach, to our knowledge not previously used in plants (Grunt *et al*. 2020), revealed similar morphology and size distributions for EVs isolated by both methods (Figure 2, Figure 3). However, there were notable differences in reproducibility, reflected by the larger confidence intervals for samples derived by ultracentrifugation in NTA (Figure 3). This is in line with PME having higher recovery rates from the used starting material, at the cost of lower specificity (Théry *et al*. 2018). Nevertheless, we consider PME a viable alternative for isolating EVs in barley and other plants, especially when larger EV quantities are needed for downstream analyses, because changes induced by *B. hordei* inoculation are still reflected with high fidelity (Figure 3). Additionally, size exclusion chromatography proved effective in further purifying barley EVs following their isolation by PME (Supplemental Figure 7).

An initial characterisation of EVs isolated from the leaf apoplast of naïve (untreated) barley plants (Schlemmer *et al*. 2021) exhibited similar morphology of vesicle-like structures as observed in our electron micrographs (Figure 2C. Moreover, the size range of EVs as estimated from electron micrographs (∼100 nm – 250 nm) reported by Schlemmer and co-workers overlaps substantially with our findings(Figure 2C; (Schlemmer *et al*. 2021)). However, we noted discrepancies in size estimates by NTA measurements between the published and our own data; these may be caused by various factors such as the barley cultivar used or differences in experimental procedures and sample handling (Figure 3; (Schlemmer *et al*. 2021)). This divergence emphasises the necessity for standardised documentation of protocols to enable comparability of findings within and between species (reviewed in (Pinedo *et al*. 2021)). In a recent report from the monocot *Sorghum bicolor*, EVs from the P40 fraction were also reported to be polydisperse with a mean size of ∼170 nm and ∼190 nm after density purification (Chaya *et al*. 2023). Notably, comparison of monocot- (∼100 nm – 500 nm, Figure 3; (Schlemmer *et al*. 2021; Chaya *et al*. 2023)) and dicot-derived EVs (∼50 nm – 300 nm, (Rutter & Innes 2017; Janda *et al*. 2023) from the present and previous reports reveals slight differences in their size distribution but could also reflect species-specific variation.

Our study suggests that the accumulation of barley EVs is an active response of the host plant to inoculation with the obligate biotrophic pathogen, *B. hordei*. We show that in particular the quantities of EVs between ∼100 nm - 300 nm in diameter increase up to ∼10- to 50-fold higher at 96 hpi and 120 hpi compared to the non-infected control sample (Figure 4). This substantial increase in EV numbers in later infection stages coincides with fungal microcolony development and incipient leaf surface proliferation (Both *et al*. 2005). Whether EV accumulation is a host response linked to particular infection stages or simply correlates with the amount of fungal biomass remains an open question. However, we argue that the altered EV populations primarily originate from the plant. This notion is supported by the fact that the predominant ∼100 nm - 300 nm EV population is already detectable (at basal levels) in the samples from non-inoculated leaves. In addition, the fungal mode of infection is restricted to the epidermal cell layer, rendering a substantial contribution to the total leaf EV population rather unlikely.

Vesicles of ∼300 nm - 500 nm in diameter exhibited similar accumulation during fungal infection to the ∼100 nm - 300 nm vesicles. However, while they also increased markedly at 96 hpi and 120 hpi, they reached considerably lower levels than the ∼100 nm - 300 nm EV population (only ∼2- to 3-fold as compared to ∼10- to 50-fold; Figure 4A). Based on this distinction we propose that (1) the larger-sized vesicle population might be different from the smaller-sized main population and that (2) these vesicles are potentially secreted *via* a different route. The latter notion is based on the fact that in other systems the biogenesis pathway is an important factor determining EV size (Mathivanan *et al*. 2010). Accordingly, the EV populations could be also associated with different cargo molecules. However, we cannot rule out that the larger vesicles of ∼300 nm - 500 nm in diameter result from artefactual fusion events of vesicles from the ∼100 nm - 300 nm population that accumulate to very high levels during fungal infection, generated during our experimental procedures.

The t-SNARE protein Ror2 is an ortholog of a known EV marker (PEN1) in plants like Arabidopsis, *Nicotiana benthamiana* and sorghum (Rutter & Innes 2017; Chaya *et al*. 2023; Zhang *et al*. 2020). We detected two specific Ror2 signals in the immunoblots of vesicle samples, corresponding to 34 kDa and ∼75 kDa in molecular mass. These signals showed an opposing dynamic in the course of fungal infection (Figure 4C). While the 34-kDa signal is indicative of monomeric Ror2 (Collins *et al*. 2003), the ∼75-kDa signal potentially represents a (homo)dimer or a Ror2-containing SNARE complex (Arien *et al*. 2003). SDS resistance of the ∼75-kDa signal rather suggests a SNARE complex, though urea resistance (Supplemental Figure 5E) also challenges this interpretation (Fasshauer *et al*. 2002; Kubista *et al*. 2004). Hence, the nature and biological relevance of the ∼75-kDa signal remains unclear at this stage. Despite its detection in EV immunoblots (Figure 4C), Ror2 was not found in our proteomics data (Table 1). This is likely because membrane-resident proteins are generally more challenging to detect, which is also reflected by the overall low number of identified proteins with transmembrane domains in our dataset (six proteins, ∼9%, Table 1), a common phenomenon in protein mass spectrometry (Souda *et al*. 2011; Bender & Schmidt 2019). On the other hand, a trypsin protection assay using the Ror2 antibody indicated its presence in EVs as cargo, making it, together with the observation of the specific ∼75-kDa signal eluting in EV fractions in SEC, a strong EV marker candidate (Figure 5). The fact that we did not detect Ror2 with the αPEN1 antibody in the detergent-only treatment indicates interference of TritonX-100 with at least the PEN1 antibody and underscores the importance of incorporating this control in protease protection assays.

EVs from Arabidopsis, sunflower, and sorghum, as well as high-speed centrifugation pellets collected from tomato cell culture medium, are associated with stress-related proteins (Rutter & Innes 2017; Regente *et al*. 2017; He *et al*. 2021; Gonorazky *et al*. 2012). Consistent with these earlier findings, the barley EV proteome was also enriched in stress-related proteins, particularly after inoculation with *B. hordei* (Table 1). However, their number in our dataset derived from crude EVs was substantially higher as compared to the proteomes of purified EVs. This discrepancy might be explained by the loose association of this type of proteins with EVs, which might get lost during purification by size exclusion chromatography. In line with the high number of stress-related proteins in crude EVs, we found two abundant pathogenesis-related proteins (PR-1 and PR-17b) that did not co-purify in size exclusion chromatography with the vesicles (Figure 5A). However, the trypsin protection assay indicated that PR-17b is protected (Figure 5C), perhaps due to interaction with glycoproteins on the EV surface. This protection might be compromised upon membrane destabilisation following detergent addition or by shear forces during size exclusion chromatography. We speculate that this behaviour indicates that PR-17b (and perhaps other (PR) proteins) are constituents of an acquired vesicle corona which forms after secretion and might play a role in EV uptake and other functions (Yerneni *et al*. 2022; Tóth *et al*. 2021; Liam-Or *et al*. 2024).

The current study dissects time-dependent EV release upon biotic stress (powdery mildew infection). Thus, it serves as a starting point for investigating the role of EVs at different stages of plant-microbe interactions. By establishing PME of vesicles, a scalable alternative method for plant EV isolation, the output of which is comparable to the routinely used ultracentrifugation, we enable feasible EV analysis in a challenging plant-microbe system, the interaction of barley with *B. hordei*. Combined with size exclusion chromatography, PME is an alternative for the isolation and purification of plant EVs. Furthermore, we unravel an infection stage-dependent EV response involving distinct EV populations, a phenomenon worth exploring in other plant-microbe interactions. Finally, our findings add to the growing evidence linking plant EVs to stress-related proteins. In conjunction with our recent data for cross-kingdom RNA interference mediated by plant EVs (Kusch *et al*. 2023), our study signifies the involvement of barley leaf EVs in the *B. hordei* interaction, prompting future investigations into the roles of the distinct EV populations.

## Acknowledgements

We thank Prof. Wilhelm Jahnen-Dechent for granting access to the NTA machine and Steffen Gräber for the great technical support. We further thank Prof. Hans-Thordal Christensen for providing PR-17 and PEN1 antibodies. We thank all lab members for fruitful discussions.

## Data availability

The mass spectrometry proteomics data have been deposited to the ProteomeXchange Consortium via the PRIDE partner repository (Perez-Riverol *et al*. 2022) with the dataset identifier PXD050823 (Reviewer login details: Username; Password: xEZRIXVN).

## Author contributions

RP, HT and PDS conceived the study. HT, CH and CK performed immunoblots. MB conducted electron microscopy. HT performed NTA and protection assays. FD executed mass spectrometry. HT analysed the data. HT drafted the manuscript. RP and PDS edited the manuscript with the help of co-author contributions. All authors have read and approved the final manuscript version.

## Abbreviations

AWF: apoplastic wash fluid
EV: extracellular vesicle
Hpi: hours post inoculation
NTA: nanoparticle tracking analysis
PME: polymer-mediated enrichment
RuBisCO: ribulose-1,5-bisphophate carboxylase/oxygenase

## References

1. Agrawal GK, Jwa N-S, Lebrun M-H, Job D, Rakwal R (2010) Plant secretome: unlocking secrets of the secreted proteins. Proteomics, doi: 10.1002/pmic.200900514.

2. Ahmed NU, Park J-I, Seo M-S et al. (2012) Identification and expression analysis of chitinase genes related to biotic stress resistance in *Brassica*. Molecular Biology Reports, doi: 10.1007/s11033-011-1139-x.

3. Aist JR, Williams PH (1971) The cytology and kinetics of cabbage root hair penetration by *Plasmodiophora brassicae*. Canadian Journal of Botany, doi: 10.1139/b71-284.

4. Almagro Armenteros JJ, Tsirigos KD, Sønderby CK et al. (2019) SignalP 5.0 improves signal peptide predictions using deep neural networks. Nature Biotechnology, doi: 10.1038/s41587-019-0036-z.

5. An QL, Hückelhoven R, Kogel KH, van Bel AJE (2006) Multivesicular bodies participate in a cell wall-associated defence response in barley leaves attacked by the pathogenic powdery mildew fungus. Cellular Microbiology, 8, 1009–1019.

6. Anisimova OK, Shchennikova AV, Kochieva EZ, Filyushin MA (2021) Pathogenesis-Related genes of PR1, PR2, PR4, and PR5 families are involved in the response to *Fusarium* infection in garlic (*Allium sativum* L.). International Journal of Molecular Sciences, doi: 10.3390/ijms22136688.

7. Arien H, Wiser O, Arkin IT, Leonov H, Atlas D (2003) Syntaxin 1A modulates the voltage-gated L-type calcium channel (Ca(v)1.2) in a cooperative manner. Journal of Biological Chemistry, doi: 10.1074/jbc.M301401200.

8. Bağcı C, Sever-Bahcekapili M, Belder N, Bennett APS, Erdener ŞE, Dalkara T (2022) Overview of extracellular vesicle characterization techniques and introduction to combined reflectance and fluorescence confocal microscopy to distinguish extracellular vesicle subpopulations. Neurophotonics, doi: 10.1117/1.NPh.9.2.021903.

9. Beers EP, Jones AM, Dickerman AW (2004) The S8 serine, C1A cysteine and A1 aspartic protease families in Arabidopsis. Phytochemistry, doi: 10.1016/j.phytochem.2003.09.005.

10. Beffa RS, Neuhaus JM, Meins F (1993) Physiological compensation in antisense transformants: specific induction of an “ersatz” glucan endo-1,3-beta-glucosidase in plants infected with necrotizing viruses. Proceedings of the National Academy of Sciences of the United States of America, doi: 10.1073/pnas.90.19.8792.

11. Bellis D de, Kalmbach L, Marhavy P, Daraspe J, Geldner N, Barberon M (2022) Extracellular vesiculo-tubular structures associated with suberin deposition in plant cell walls. Nature Communications, doi: 10.1038/s41467-022-29110-0.

12. Bender J, Schmidt C (2019) Mass spectrometry of membrane protein complexes. Biological Chemistry, doi: 10.1515/hsz-2018-0443.

13. Boersema PJ, Raijmakers R, Lemeer S, Mohammed S, Heck AJR (2009) Multiplex peptide stable isotope dimethyl labeling for quantitative proteomics. Nature Protocols, doi: 10.1038/nprot.2009.21.

14. Both M, Csukai M, Stumpf MPH, Spanu PD (2005) Gene expression profiles of *Blumeria graminis* indicate dynamic changes to primary metabolism during development of an obligate biotrophic pathogen. Plant Cell, 17, 2107–2122.

15. Bowien B, Mayer F (1978) Further studies on the quaternary structure of D-ribulose-1, 5- bisphosphate carboxylase from *Alcaligenes eutrophus*. European Journal of Biochemistry, doi: 10.1111/j.1432-1033.1978.tb12426.x.

16. Bryngelsson T, Sommer-Knudsen J, Gregersen PL, Collinge DB, Ek B, Thordal-Christensen H (1994) Purification, characterization, and molecular cloning of basic PR-1-type pathogenesis-related proteins from barley. Molecular Plant-Microbe Interactions, doi: 10.1094/MPMI-7-0267.

17. Cai Q, He B, Jin H (2019) A safe ride in extracellular vesicles - small RNA trafficking between plant hosts and pathogens. Current Opinion in Plant Biology, doi: 10.1016/j.pbi.2019.09.001.

18. Cai Q, He B, Wang S et al. (2021) Message in a bubble: Shuttling small RNAs and proteins between cells and interacting organisms using extracellular vesicles. Annual Review of Plant Biology, doi: 10.1146/annurev-arplant-081720-010616.

19. Cai Q, Qiao L, Wang M et al. (2018) Plants send small RNAs in extracellular vesicles to fungal pathogen to silence virulence genes. Science, doi: 10.1126/science.aar4142.

20. Caruso Bavisotto C, Cappello F, Macario AJL et al. (2017) Exosomal HSP60: a potentially useful biomarker for diagnosis, assessing prognosis, and monitoring response to treatment. Expert Review of Molecular Diagnostics, doi: 10.1080/14737159.2017.1356230.

21. Chaya T, Banerjee A, Rutter BD et al. (2023) The extracellular vesicle proteomes of *Sorghum bicolor* and *Arabidopsis thaliana* are partially conserved. Plant Physiology, doi: 10.1093/plphys/kiad644.

22. Chernyshev VS, Rachamadugu R, Tseng YH et al. (2015) Size and shape characterization of hydrated and desiccated exosomes. Analytical and Bioanalytical Chemistry, doi: 10.1007/s00216-015-8535-3.

23. Chevallet M, Luche S, Rabilloud T (2006) Silver staining of proteins in polyacrylamide gels. Nature Protocols, doi: 10.1038/nprot.2006.288.

24. Choi D-S, Kim D-K, Kim Y-K, Gho YS (2015) Proteomics of extracellular vesicles: Exosomes and ectosomes. Mass Spectrometry Reviews, doi: 10.1002/mas.21420.

25. Chow FW-N, Koutsovoulos G, Ovando-Vázquez C et al. (2019) Secretion of an Argonaute protein by a parasitic nematode and the evolution of its siRNA guides. Nucleic Acids Research, doi: 10.1093/nar/gkz142.

26. Christensen AB, Cho BH, Næsby M et al. (2002) The molecular characterization of two barley proteins establishes the novel PR-17 family of pathogenesis-related proteins. Molecular Plant Pathology, doi: 10.1046/j.1364-3703.2002.00105.x.

27. Clos-Sansalvador M, Monguió-Tortajada M, Roura S, Franquesa M, Borràs FE (2022) Commonly used methods for extracellular vesicles’ enrichment: Implications in downstream analyses and use. European Journal of Cell Biology, doi: 10.1016/j.ejcb.2022.151227.

28. Cocucci E, Meldolesi J (2015) Ectosomes and exosomes: shedding the confusion between extracellular vesicles. Trends in Cell Biology, doi: 10.1016/j.tcb.2015.01.004.

29. Collins NC, Thordal-Christensen H, Lipka V et al. (2003) SNARE-protein-mediated disease resistance at the plant cell wall. Nature, 425, 973–977.

30. Colombo M, Raposo G, Théry C (2014) Biogenesis, secretion, and intercellular interactions of exosomes and other extracellular vesicles. Annual Review of Cell and Developmental Biology, doi: 10.1146/annurev-cellbio-101512-122326.

31. Dmochowska-Boguta M, Nadolska-Orczyk A, Orczyk W (2013) Roles of peroxidases and NADPH oxidases in the oxidative response of wheat (*Triticum aestivum*) to brown rust (*Puccinia triticina*) infection. Plant Pathology, doi: 10.1111/ppa.12009.

32. Doehlemann G, Hemetsberger C (2013) Apoplastic immunity and its suppression by filamentous plant pathogens. The New phytologist, doi: 10.1111/nph.12277.

33. Dos Santos C, Franco OL (2023) Pathogenesis-Related proteins (PRs) with enzyme activity activating plant defense responses. Plants, doi: 10.3390/plants12112226.

34. Du Y, Stegmann M, Misas Villamil JC (2016) The apoplast as battleground for plant-microbe interactions. New Phytologist, doi: 10.1111/nph.13777.

35. Fasshauer D, Antonin W, Subramaniam V, Jahn R (2002) SNARE assembly and disassembly exhibit a pronounced hysteresis. Nature Structural Biology, doi: 10.1038/nsb750.

36. Felle HH, Herrmann A, Hanstein S, Hückelhoven R, Kogel K-H (2004) Apoplastic pH signaling in barley leaves attacked by the powdery mildew fungus *Blumeria graminis* f. sp. *hordei*. Molecular Plant-Microbe Interactions, doi: 10.1094/MPMI.2004.17.1.118.

37. Finnie C, Andersen CH, Borch J et al. (2002) Do 14-3-3 proteins and plasma membrane H^+^- AtPases interact in the barley epidermis in response to the barley powdery mildew fungus? Plant Molecular Biology, doi: 10.1023/a:1014938417267.

38. Gardiner C, Di Vizio D, Sahoo S et al. (2016) Techniques used for the isolation and characterization of extracellular vesicles: Results of a worldwide survey. Journal of Extracellular Vesicles, doi: 10.3402/jev.v5.32945.

39. Glawe DA (2008) The powdery mildews: A review of the world’s most familiar (yet poorly known) plant pathogens. Annual Review of Phytopathology, doi: 10.1146/annurev.phyto.46.081407.104740.

40. Gonorazky G, Laxalt AM, Dekker HL et al. (2012) Phosphatidylinositol 4-phosphate is associated to extracellular lipoproteic fractions and is detected in tomato apoplastic fluids. Plant Biology, doi: 10.1111/j.1438-8677.2011.00488.x.

41. Greening DW, Simpson RJ (2018) Understanding extracellular vesicle diversity - current status. Expert review of Proteomics, doi: 10.1080/14789450.2018.1537788.

42. Grunt M, Failla AV, Stevic I, Hillebrand T, Schwarzenbach H (2020) A novel assay for exosomal and cell-free miRNA isolation and quantification. RNA Biology, doi: 10.1080/15476286.2020.1721204.

43. Halperin W, Jensen WA (1967) Ultrastructural changes during growth and embryogenesis in carrot cell cultures. Journal of Ultrastructure Research, doi: 10.1016/S0022-5320(67)80128-X.

44. He B, Cai Q, Qiao L et al. (2021) RNA-binding proteins contribute to small RNA loading in plant extracellular vesicles. Nature Plants, doi: 10.1038/s41477-021-00863-8.

45. Herrmann IK, Wood MJA, Fuhrmann G (2021) Extracellular vesicles as a next-generation drug delivery platform. Nature Nanotechnology, doi: 10.1038/s41565-021-00931-2.

46. Hill EH, Solomon PS (2020) Extracellular vesicles from the apoplastic fungal wheat pathogen *Zymoseptoria tritici*. Fungal Biology and Biotechnology, doi:10.1186/s40694-020-00103-2.

47. Hughes CS, Moggridge S, Müller T, Sorensen PH, Morin GB, Krijgsveld J (2019) Single-pot, solid-phase-enhanced sample preparation for proteomics experiments. Nature Protocols, doi: 10.1038/s41596-018-0082-x.

48. Hunt M, Banerjee S, Surana P et al. (2019) Small RNA discovery in the interaction between barley and the powdery mildew pathogen. BMC Genomics, *doi:* 10.1186/s12864-019-5947-z.

49. Huokko T, Ni T, Dykes GF et al. (2021) Probing the biogenesis pathway and dynamics of thylakoid membranes. Nature Communications, doi: 10.1038/s41467-021-23680-1.

50. Hurwitz SN, Rider MA, Bundy JL, Liu X, Singh RK, Meckes DG (2016) Proteomic profiling of NCI-60 extracellular vesicles uncovers common protein cargo and cancer type-specific biomarkers. Oncotarget, doi: 10.18632/oncotarget.13569.

51. Hwang S-K, Singh S, Cakir B, Satoh H, Okita TW (2016) The plastidial starch phosphorylase from rice endosperm: Catalytic properties at low temperature. Planta, doi: 10.1007/s00425-015-2461-7.

52. Hyun TK, van der Graaff E, Albacete A et al. (2014) The *Arabidopsis* PLAT domain protein1 is critically involved in abiotic stress tolerance. PLoS One, doi: 10.1371/journal.pone.0112946.

53. Janda M, Rybak K, Krassini L et al. (2023) Biophysical and proteomic analyses of *Pseudomonas syringae* pv. *tomato* DC3000 extracellular vesicles suggest adaptive functions during plant infection. mBio, doi: 10.1128/mbio.03589-22.

54. Jashni MK, Mehrabi R, Collemare J, Mesarich CH, Wit PJGM de (2015) The battle in the apoplast: Further insights into the roles of proteases and their inhibitors in plant- pathogen interactions. Frontiers in Plant Science, doi: 10.3389/fpls.2015.00584.

55. Ju S, Mu J, Dokland T et al. (2013) Grape exosome-like nanoparticles induce intestinal stem cells and protect mice from DSS-induced colitis. Molecular Therapy, doi: 10.1038/mt.2013.64.

56. Jumper J, Evans R, Pritzel A et al. (2021) Highly accurate protein structure prediction with AlphaFold. Nature, doi: 10.1038/s41586-021-03819-2.

57. Kayum MA, Park J-I, Nath UK, Biswas MK, Kim H-T, Nou I-S (2017) Genome-wide expression profiling of aquaporin genes confer responses to abiotic and biotic stresses in *Brassica rapa*. BMC Plant Biology, doi: 10.1186/s12870-017-0979-5.

58. Krogh A, Larsson B, Heijne G von, Sonnhammer EL (2001) Predicting transmembrane protein topology with a hidden Markov model: application to complete genomes. Journal of Molecular Biology, doi: 10.1006/jmbi.2000.4315.

59. Kubista H, Edelbauer H, Boehm S (2004) Evidence for structural and functional diversity among SDS-resistant SNARE complexes in neuroendocrine cells. Journal of Cell Science, doi: 10.1242/jcs.00941.

60. Kusch S, Frantzeskakis L, Thieron H, Panstruga R (2018) Small RNAs from cereal powdery mildew pathogens may target host plant genes. Fungal Biology, doi: 10.1016/j.funbio.2018.08.008.

61. Kusch S, Singh M, Thieron H, Spanu PD, Panstruga R (2023) Site-specific analysis reveals candidate cross-kingdom small RNAs, tRNA and rRNA fragments, and signs of fungal RNA phasing in the barley-powdery mildew interaction. Molecular Plant Pathology, doi: 10.1111/mpp.13324.

62. Kusumawati L, Imin N, Djordjevic MA (2008) Characterization of the secretome of suspension cultures of *Medicago* species reveals proteins important for defense and development. Journal of Proteome Research, doi: 10.1021/pr800291z.

63. Kwon S, Rupp O, Brachmann A et al. (2021) mRNA inventory of extracellular vesicles from *Ustilago maydis*. Journal of Fungi, doi: 10.3390/jof7070562.

64. La Canal L de, Pinedo M (2018) Extracellular vesicles: A missing component in plant cell wall remodeling. Journal of Experimental Botany, doi: 10.1093/jxb/ery255.

65. Lambertucci S, Orman KM, Das Gupta S et al. (2019) Analysis of barley leaf epidermis and extrahaustorial proteomes during powdery mildew infection reveals that the PR5 thaumatin-like protein TLP5 Is required for susceptibility towards *Blumeria graminis* f. sp. *hordei*. Frontiers in Plant Science, doi: 10.3389/fpls.2019.01138.

66. Li D, Li Z, Wu J et al. (2022) Analysis of outer membrane vesicles indicates that glycerophospholipid metabolism contributes to early symbiosis between *Sinorhizobium fredii* HH103 and soybean. Molecular Plant-Microbe Interactions, *doi:* 10.1094/MPMI-11-21-0288-R.

67. Liam-Or R, Faruqu FN, Walters A et al. (2024) Cellular uptake and in vivo distribution of mesenchymal-stem-cell-derived extracellular vesicles are protein corona dependent. Nature Nanotechnology, doi: 10.1038/s41565-023-01585-y.

68. Lu P-P, Yu T-F, Zheng W-J et al. (2018) The wheat Bax inhibitor-1 protein interacts with an aquaporin TaPIP1 and enhances disease resistance in *Arabidopsis*. Frontiers in Plant Science, doi: 10.3389/fpls.2018.00020.

69. Martínez-Greene JA, Hernández-Ortega K, Quiroz-Baez R et al. (2021) Quantitative proteomic analysis of extracellular vesicle subgroups isolated by an optimized method combining polymer-based precipitation and size exclusion chromatography. Journal of Extracellular Vesicles, doi: 10.1002/jev2.12087.

70. Mathivanan S, Ji H, Simpson RJ (2010) Exosomes: Extracellular organelles important in intercellular communication. Journal of Proteomics, doi: 10.1016/j.jprot.2010.06.006.

71. Micali CO, Neumann U, Grunewald D, Panstruga R, O’Connell R (2011) Biogenesis of a specialized plant-fungal interface during host cell internalization of *Golovinomyces orontii* haustoria. Cellular Microbiology, 13, 210–226.

72. Misas Villamil JC, Mueller AN, Demir F et al. (2019) A fungal substrate mimicking molecule suppresses plant immunity via an inter-kingdom conserved motif. Nature Communications, doi: 10.1038/s41467-019-09472-8.

73. Misas-Villamil JC, van der Hoorn RAL, Doehlemann G (2016) Papain-like cysteine proteases as hubs in plant immunity. New Phytologist, doi: 10.1111/nph.14117.

74. Mostek A, Börner A, Badowiec A, Weidner S (2015) Alterations in root proteome of salt- sensitive and tolerant barley lines under salt stress conditions. Journal of Plant Physiology, doi: 10.1016/j.jplph.2014.08.020.

75. Mulaosmanovic E, Lindblom TUT, Bengtsson M et al. (2020) High-throughput method for detection and quantification of lesions on leaf scale based on trypan blue staining and digital image analysis. Plant Methods, doi: 10.1186/s13007-020-00605-5.

76. Nasfi S, Kogel K-H (2022) Packaged or unpackaged: Appearance and transport of extracellular noncoding RNAs in the plant apoplast. ExRNA, doi: 10.21037/exrna-22-11.

77. Nowara D, Gay A, Lacomme C et al. (2010) HIGS: Host-Induced Gene Silencing in the obligate biotrophic fungal pathogen *Blumeria graminis*. Plant Cell, 22, 3130–3141.

78. Pál M, Kovács V, Vida G, Szalai G, Janda T (2013) Changes induced by powdery mildew in the salicylic acid and polyamine contents and the antioxidant enzyme activities of wheat lines. European Journal of Plant Pathology, doi: 10.1007/s10658-012-0063-9.

79. Panstruga R, Spanu P (2024) Transfer RNA and ribosomal RNA fragments - emerging players in plant-microbe interactions. New Phytologist, doi: 10.1111/nph.19409.

80. Pečenková T, Pejchar P, Moravec T et al. (2022) Immunity functions of Arabidopsis pathogenesis-related 1 are coupled but not confined to its C-terminus processing and trafficking. Molecular Plant Pathology, doi: 10.1111/mpp.13187.

81. Perez-Riverol Y, Bai J, Bandla C et al. (2022) The PRIDE database resources in 2022: A hub for mass spectrometry-based proteomics evidences. Nucleic Acids Research, doi: 10.1093/nar/gkab1038.

82. Pinedo M, La Canal L de, Marcos Lousa C de (2021) A call for Rigor and standardization in plant extracellular vesicle research. Journal of Extracellular Vesicles, doi: 10.1002/jev2.12048.

83. Prado N, Alché JdD, Casado-Vela J et al. (2014) Nanovesicles are secreted during pollen germination and pollen tube growth: A possible role in fertilization. Molecular Plant, doi: 10.1093/mp/sst153.

84. Qiao SA, Gao Z, Roth R (2023) A perspective on cross-kingdom RNA interference in mutualistic symbioses. New Phytologist, doi: 10.1111/nph.19122.

85. Raunser S, Magnani R, Huang Z et al. (2009) Rubisco in complex with Rubisco large subunit methyltransferase. Proceedings of the National Academy of Sciences of the United States of America, doi: 10.1073/pnas.0810563106.

86. Regente M, Corti-Monzón G, Maldonado AM, Pinedo M, Jorrín J, La Canal L de (2009) Vesicular fractions of sunflower apoplastic fluids are associated with potential exosome marker proteins. FEBS Letters, doi: 10.1016/j.febslet.2009.09.041.

87. Regente M, Pinedo M, San Clemente H, Balliau T, Jamet E, La Canal L de (2017) Plant extracellular vesicles are incorporated by a fungal pathogen and inhibit its growth. Journal of Experimental Botany, doi: 10.1093/jxb/erx355.

88. Rohringer R, Ebrahim-Nesbat F, Wolf G (1983) Proteins in intercellular washing fluids from leaves of barley (*Hordeum vulgare* L.). Journal of Experimental Botany, doi: 10.1093/jxb/34.12.1589.

89. Roth R, Hillmer S, Funaya C et al. (2019) Arbuscular cell invasion coincides with extracellular vesicles and membrane tubules. Nature Plants, doi: 10.1038/s41477-019-0365-4.

90. Ruf A, Oberkofler L, Robatzek S, Weiberg A (2022) Spotlight on plant RNA-containing extracellular vesicles. Current Opinion in Plant Biology, doi: 10.1016/j.pbi.2022.102272.

91. Rutter BD, Chu T-T-H, Dallery J-F, Zajt KK, O’Connell RJ, Innes RW (2022) The development of extracellular vesicle markers for the fungal phytopathogen *Colletotrichum higginsianum*. Journal of Extracellular Vesicles, doi: 10.1002/jev2.12216.

92. Rutter BD, Innes RW (2017) Extracellular vesicles isolated from the leaf apoplast carry stress- response proteins. Plant Physiology, doi: 10.1104/pp.16.01253.

93. Rutter BD, Rutter KL, Innes RW (2017) Isolation and quantification of plant extracellular vesicles. Bio-protocol, doi: 10.21769/BioProtoc.2533.

94. Rybak K, Robatzek S (2019) Functions of extracellular vesicles in immunity and virulence. Plant Physiology, doi: 10.1104/pp.18.01557.

95. Santén K, Marttila S, Liljeroth E, Bryngelsson T (2005) Immunocytochemical localization of the pathogenesis-related PR-1 protein in barley leaves after infection by Bipolaris sorokiniana. Physiological and Molecular Plant Pathology, doi: 10.1016/j.pmpp.2005.04.006.

96. Sargent C (1977) Barley epidermal apoplast structure and modification by powdery mildew contact. Physiological Plant Pathology, doi: 10.1016/0048-4059(77)90058-3.

97. Schlemmer T, Barth P, Weipert L et al. (2021) Isolation and characterization of barley (*Hordeum vulgare*) extracellular vesicles to assess their role in RNA spray-based crop protection. International Journal of Molecular Sciences, doi: 10.3390/ijms22137212.

98. Segarra CI, Casalongué CA, Pinedo ML, Ronchi VP, Conde RD (2003) A germin-like protein of wheat leaf apoplast inhibits serine proteases. Journal of Experimental Botany, doi: 10.1093/jxb/erg139.

99. Shaw M, Manocha MS (1965) Fine structure in detached, senescing wheat leaves. Canadian Journal of Botany, doi: 10.1139/b65-084.

100. Solé M, Scheibner F, Hoffmeister A-K et al. (2015) *Xanthomonas campestris* pv. *vesicatoria s*ecretes proteases and xylanases via the Xps type II secretion system and outer membrane vesicles. Journal of Bacteriology, doi: 10.1128/jb.00322-15.

101. Souda P, Ryan CM, Cramer WA, Whitelegge J (2011) Profiling of integral membrane proteins and their post translational modifications using high-resolution mass spectrometry. Methods, doi: 10.1016/j.ymeth.2011.09.019.

102. Stranford DM, Leonard JN (2017) Delivery of biomolecules via extracellular vesicles: A budding therapeutic strategy. Advances in Genetics, doi: 10.1016/bs.adgen.2017.08.002.

103. Tamás L, Huttová J, Žigová Z (1997) Accumulation of stress-proteins in intercellular spaces of barley leaves induced by biotic and abiotic factors. Biologia Plantarum, doi: 10.1023/A:1001028226434.

104. Taylor DD, Shah S (2015) Methods of isolating extracellular vesicles impact down-stream analyses of their cargoes. Methods, doi: 10.1016/j.ymeth.2015.02.019.

105. Théry C, Amigorena S, Raposo G, Clayton A (2006) Isolation and characterization of exosomes from cell culture supernatants and biological fluids. Current Protocols, doi: 10.1002/0471143030.cb0322s30.

106. Théry C, Witwer KW, Aikawa E et al. (2018) Minimal information for studies of extracellular vesicles 2018 (MISEV2018): A position statement of the International Society for Extracellular Vesicles and update of the MISEV2014 guidelines. Journal of Extracellular Vesicles, doi: 10.1080/20013078.2018.1535750.

107. Thomas M, Huck N, Hoehenwarter W, Conrath U, Beckers GJM (2015) Combining metabolic ^15^N labeling with improved tandem MOAC for enhanced probing of the phosphoproteome. In: Plant Phosphoproteomics. (ed. Schulze WX), pp. 81–96. Springer New York, New York, NY.

108. Tkach M, Théry C (2016) Communication by extracellular vesicles: Where we are and where we need to go. Cell, doi: 10.1016/j.cell.2016.01.043.

109. Tóth EÁ, Turiák L, Visnovitz T et al. (2021) Formation of a protein corona on the surface of extracellular vesicles in blood plasma. Journal of Extracellular Vesicles, doi: 10.1002/jev2.12140.

110. Tyanova S, Temu T, Cox J (2016) The MaxQuant computational platform for mass spectrometry-based shotgun proteomics. Nature Protocols, doi: 10.1038/nprot.2016.136.

111. U Stotz H, Brotherton D, Inal J (2022) Communication is key: Extracellular vesicles as mediators of infection and defence during host-microbe interactions in animals and plants. FEMS Microbiology Reviews, doi: 10.1093/femsre/fuab044.

112. Vogel R, Savage J, Muzard J et al. (2021) Measuring particle concentration of multimodal synthetic reference materials and extracellular vesicles with orthogonal techniques: Who is up to the challenge? Journal of Extracellular Vesicles, doi: 10.1002/jev2.12052.

113. Wang J, Ding Y, Wang J et al. (2010) EXPO, an exocyst-positive organelle distinct from multivesicular endosomes and autophagosomes, mediates cytosol to cell wall exocytosis in Arabidopsis and tobacco cells. Plant Cell, doi: 10.1105/tpc.110.080697.

114. Wang L, Fobert PR (2013) Arabidopsis clade I TGA factors regulate apoplastic defences against the bacterial pathogen *Pseudomonas syringae* through endoplasmic reticulum- based processes. PLoS One, doi: 10.1371/journal.pone.0077378.

115. Wang S, He B, Wu H et al. (2024) Plant mRNAs move into a fungal pathogen via extracellular vesicles to reduce infection. Cell Host & Microbe, doi: 10.1016/j.chom.2023.11.020.

116. Yerneni SS, Solomon T, Smith J, Campbell PG (2022) Radioiodination of extravesicular surface constituents to study the biocorona, cell trafficking and storage stability of extracellular vesicles. Biochimica Et Biophysica Acta, doi: 10.1016/j.bbagen.2021.130069.

117. Yu K, Wei L, Yuan H et al. (2022) Genetic architecture of inducible and constitutive metabolic profile related to drought resistance in qingke (Tibetan hulless barley). Frontiers in Plant Science, doi: 10.3389/fpls.2022.1076000.

118. Zand Karimi H, Baldrich P, Rutter BD et al. (2022) Arabidopsis apoplastic fluid contains sRNA- and circular RNA-protein complexes that are located outside extracellular vesicles. Plant Cell, doi: 10.1093/plcell/koac043.

119. Zhang J, Qiu Y, Xu K (2020) Characterization of GFP-AtPEN1 as a marker protein for extracellular vesicles isolated from *Nicotiana benthamiana* leaves. Plant Signaling & Behavior, doi: 10.1080/15592324.2020.1791519.

120. Zhao T-Y, Corum III JW, Mullen J et al. (2006) An alkaline α-galactosidase transcript is present in maize seeds and cultured embryo cells, and accumulates during stress. Seed Science Research, doi: 10.1079/SSR2006243.

121. Zierold U, Scholz U, Schweizer P (2005) Transcriptome analysis of *mlo*-mediated resistance in the epidermis of barley. Molecular Plant Pathology, doi:10.1111/j.1364-3703.2005.00271.x.

