## Supplemental Figures for "Barley powdery mildew invasion coincides with the dynamic accumulation of leaf apoplastic extracellular vesicles that are associated with host stress response proteins"

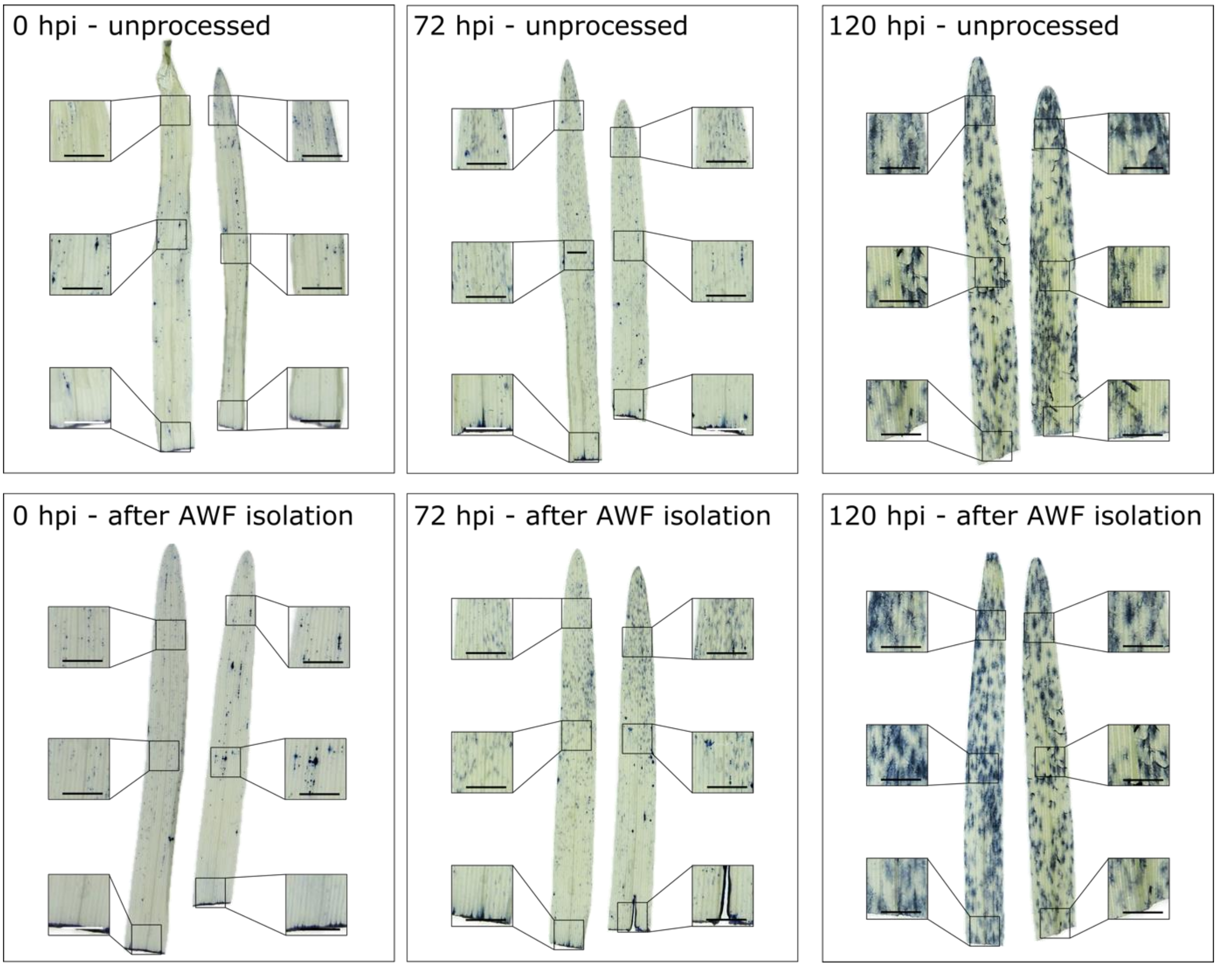


**Supplemental Figure 1. AWF isolation does not considerably affect the integrity of plant cells.** Leaves of barley plants were harvested either prior to inoculation (0 hpi) or at 24 hpi, 72 hpi, or 120 hpi with *B. hordei* and either used for isolation of AWF or left unprocessed (no buffer infiltration). After removal of leaf pigments, dead cells and fungal hyphae were stained by trypan blue. Two representative leaves with six representative regions (magnifications in boxes) per time point and treatment are shown. Scale bars = 0.5 cm.

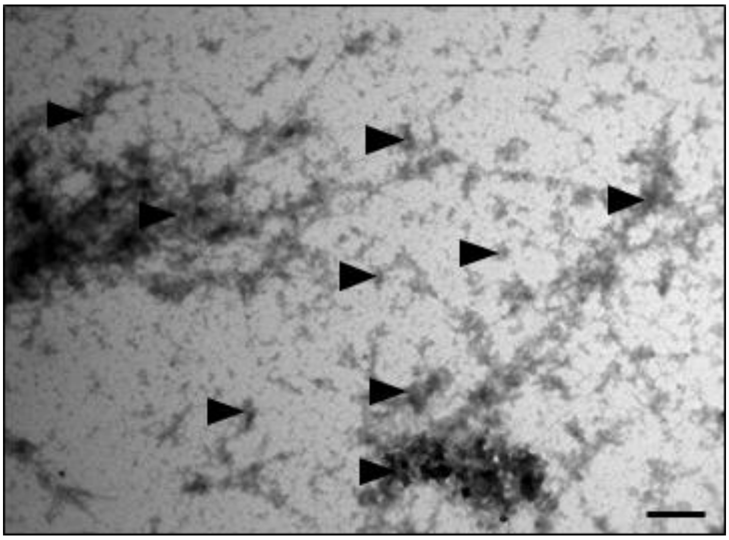

**Supplemental Figure 2. Uranyl acetate stains mannuronate/guluronate polymers used for PME of AWF-derived EVs**. Transmission electron micrographs of EV fractions isolated via PME and stained with uranyl acetate. Stained mannuronate/guluronate polymers are marked with black arrowheads. Scale bar = 1 µm.

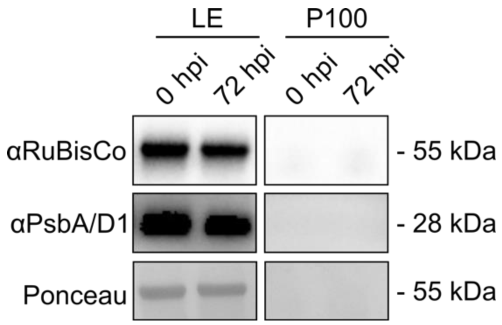


**Supplemental Figure 3. RuBisCO is undetectable in P100 fractions derived from barley leaf AWF.** Immunoblot of crude leaf extract (left panel) and the P100 fraction (right panel) probed with αRuBisCO and αPsbA/D1 (markers for chloroplastic/cytosolic contamination) antibodies. Leaf extract, and AWF-derived P100 fractions were isolated from leaves of 10-day-old barley plants that were sampled either prior to inoculation (0 hpi) or at 72 hpi with *B. hordei*. Staining with Ponceau S (the prominent band corresponding to the large subunit of RuBisCO) served to demonstrate equal loading. Apparent molecular masses of proteins (given on the right) were derived from a comparison with molecular mass standards analysed on the same gel. The experiment was repeated in two independent biological replicates with similar results.

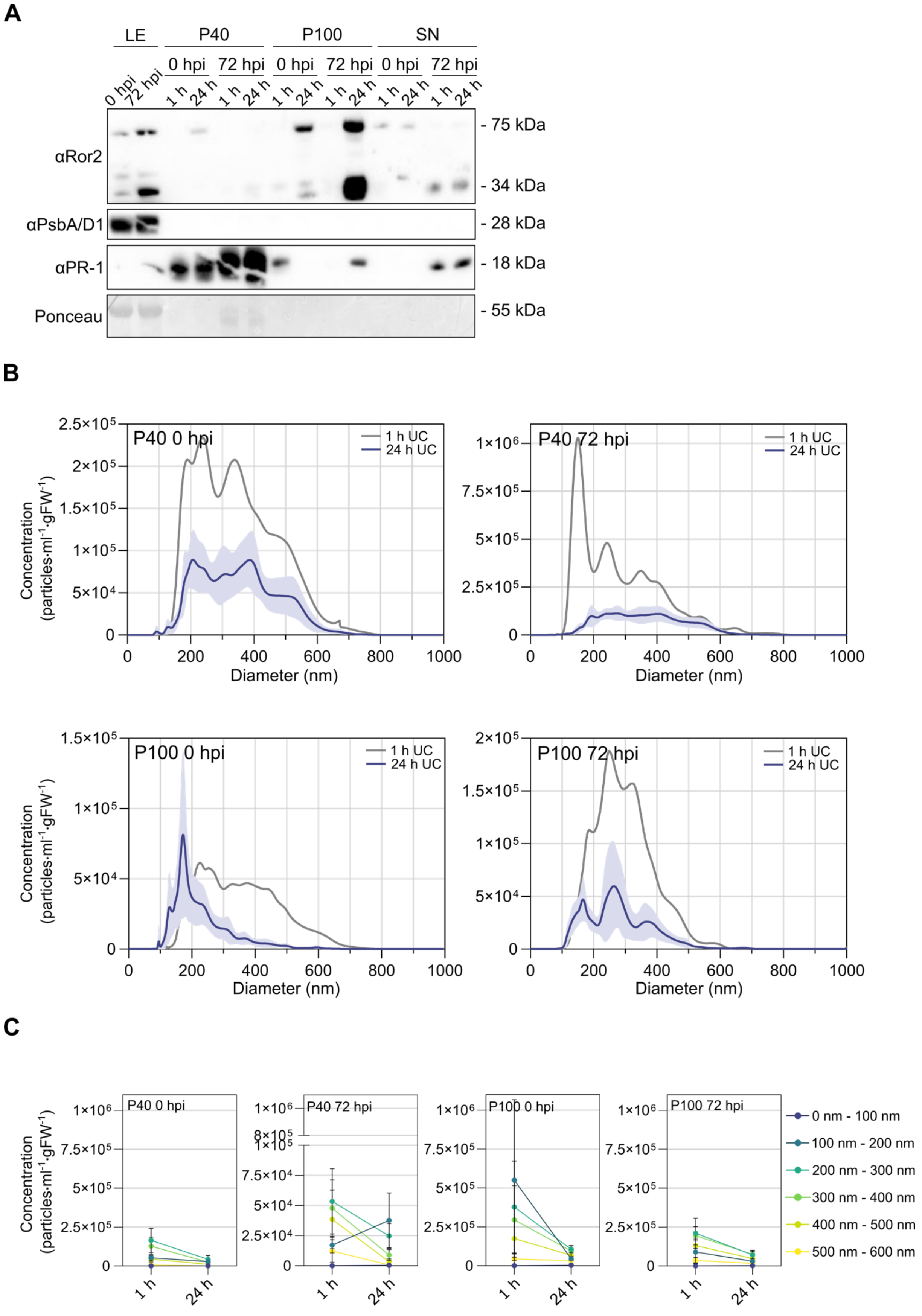
A
LE
P40
P100
SN
0 hpi
72 hpi
1 h
24 h
αRor2
αPsbA/D1
αPR-1
Ponceau
- 75 kDa
- 34 kDa
- 28 kDa
- 18 kDa
- 55 kDa
B
P40 0 hpi
P40 72 hpi
P100 0 hpi
P100 72 hpi
1 h UC
24 h UC
Concentration (particles·ml-1·gFW-1)
Diameter (nm)
C
0 nm - 100 nm
100 nm - 200 nm
200 nm - 300 nm
300 nm - 400 nm
400 nm - 500 nm
500 nm - 600 nm

**Supplemental Figure 4. Characteristics of EVs isolated with 1 h or 24 h of high-speed centrifugation.**

**A** Immunoblot probed with antibodies specific for the potential EV marker Ror2 (predicted molecular mass 34 kDa), the chloroplastic/cytosolic contamination marker PsbA/D1 (predicted molecular mass 28 kDa), and the secreted defence marker PR-1 (predicted molecular mass 18 kDa). Leaf extract (LE), EV fractions (P40- and P100-derived) and supernatant (SN) were isolated from leaves of 10-day-old plants that were either sampled prior to inoculation (0 hpi) or at 72 hpi with *B. hordei* by either 1 h or 24 h centrifugation. The gel was loaded with completely resuspended EV pellets, 10 µl of the supernatant and 5 µg leaf extract proteins. Molecular masses of proteins (given on the right) were judged according to a molecular mass standard run on the same gel. The experiment was performed independently three times with similar results.

**B** NTA data of the size distribution and mean concentration within the upper and lower 95% confidence intervals of P40- and P100-derived EV fractions isolated by 24 h of centrifugation. Shown in grey is the size distribution and mean concentration of EVs isolated from the same starting material but with 1 h of centrifugation instead. EVs were isolated from leaves of 10-day-old barley plants that were either sampled prior to inoculation (0 hpi) or at 72 hpi with *B. hordei*.

**C** Comparison of EV concentrations in EV samples isolated by either 1 h or 24 h of centrifugation, divided into stepwise 100 nm size fractions as indicated by the color-coded legend on the right side of the plots. Points represent the mean, and error bars the standard deviation. Plots are based on data from four biological replicates each.

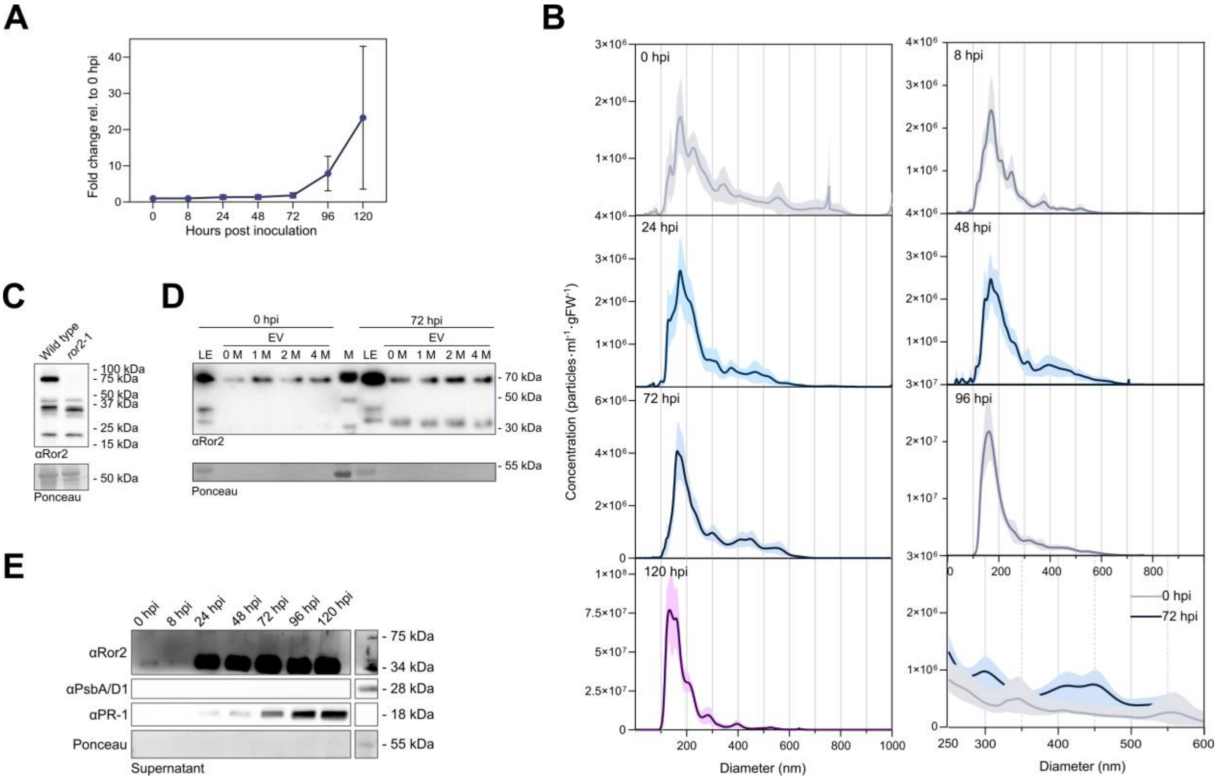


**Supplemental Figure 5. Time course analysis of EVs isolated by PME from barley leaves at 0, 8, 24, 48, 72, 96 or 120 hpi with *B. hordei*.**

**A** Total particle concentration at different time points after inoculation with *B. hordei*, relative to the particle concentration at 0 hpi. Points represent the mean, error bars standard deviation. A two-sided unpaired Welch's t-test did not reveal any statistically significant differences between time points. The plot is based on the dataset shown in Figure 4A.

**B** NTA data showing the size distribution and mean concentration within the upper and lower 95% confidence intervals of PME-derived EVs isolated from leaves of 10-day-old barley plants at different time points after inoculation with *B. hordei*. The panel in the right bottom section shows the overlay of size profiles at 0 hpi and 72 hpi in the size range of 250 nm to 600 nm. The plot and the color-coding is based on the dataset shown in Figure 4A.

**C** Immunoblots probed with antibodies specific for the potential EV marker protein Ror2 (predicted molecular mass 34 kDa) in total leaf extracts of non-inoculated wild type (*Ror2* genotype) and *ror*2-1 mutant plants. Gels were loaded with 7.5 µg of total protein extract mixed with 6x loading dye. Staining with Ponceau S served to demonstrate equal loading. The experiment was repeated independently two times with similar results.

**D** Immunoblot probed with an antibody for the potential EV marker protein Ror2 (predicted molecular mass 34 kDa) in either total leaf extract (LE) or EV samples collected either prior to inoculation (0 hpi) or at 72 hpi with *B. hordei*. EV samples were resuspended in buffer with urea concentrations ranging from 0 M - 4 M. The gel was loaded with 1.73 µg protein in 20 µl total volume with 6x loading dye per sample. Ponceau S was used as loading control for LE samples. The experiment was repeated independently two times with similar results.

**E** Immunoblots probed with antibodies specific for the potential EV marker protein Ror2 (predicted molecular mass 34 kDa), the secreted defence marker PR-1 (predicted molecular mass 18 kDa), and the chloroplastic/cytosolic contamination marker PsbA (predicted molecular mass 28 kDa) in the supernatant fraction at different time points after inoculation with *B. hordei*. Gels were loaded with 10 µl of the supernatant mixed with 6x loading dye. Molecular masses of proteins (given on the right) were estimated from comparison to a molecular mass standard analysed on the same gel. The experiment was performed in six independent biological replicates for samples collected at 0 hpi to 72 hpi and three independent biological replicates for samples collected at 96 hpi and 120 hpi.

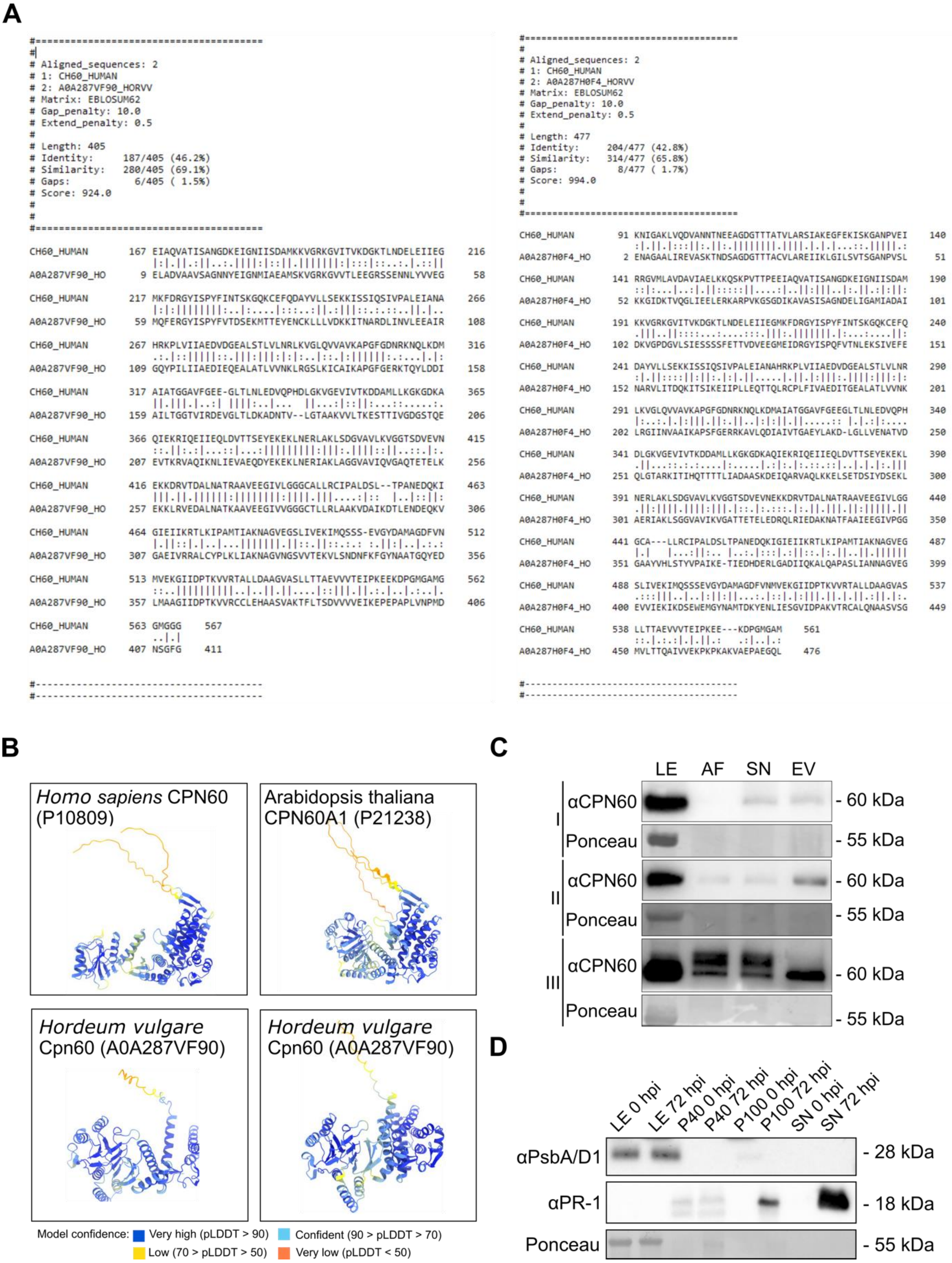


**Supplemental Figure 6. Barley CPN60 homologs might qualify as EV markers.**

**A** Pairwise alignment of human CPN60 (P10809) with barley Cpn60 A0A287VF90 and A0A287H0F4 in EMBOSS water (https://www.ebi.ac.uk/Tools/psa/emboss_water/) using the Smith-Waterman algorithm and default settings.

**B** Predicted 3D structures of *H. sapiens* CPN60 (P10809), *A. thaliana* CPN60A1 (P21238), and *H. vulgare* Cpn60s (A0A287VF90 and A0A287H0F4). Structures were predicted by AlphaFold

(Jumper *et al.* 2021) and visualised in ChimeraX (https://www.rbvi.ucsf.edu/chimerax/). AlphaFold produces a per-residue confidence (pLDDT) between 0 and 100, reflected in the model, see confidence scale.

**C** Immunoblots probed with antibodies against the potential EV marker protein CPN60 (predicted molecular mass 60 kDa) in total leaf extract (LE), AWF, P100 supernatant (SN) or PME-derived EVs isolated from leaves of 10-day-old barley plants that were sampled at 72 hpi with *B. hordei*. The gel was loaded with 5 µg protein total leaf extract (LE) or 10 µl of AWF, supernatant (SN), or PME-derived EVs mixed with 6x loading dye. Molecular masses of proteins (given on the right) were judged according to a molecular mass standard analysed on the same gel. The experiment was performed three times (I, II, III).

**D** Immunoblots probed with antibodies specific for the chloroplastic/cytosolic marker PsbA (predicted molecular mass 28 kDa) and the secreted defence marker PR-1 (predicted molecular mass 18 kDa) and in total leaf extract (LE), ultracentrifugation-derived EVs (P40 and P100) or P100 supernatant (SN) isolated from leaves of 10-day-old barley plants that were sampled either prior to inoculation (0 hpi) or at 72 hpi with *B. hordei*. The gels was loaded with 20 µl of EV pellets resuspended in Tris buffer (pH 7.5) and mixed with 6x loading dye. For leaf extract, 5 µg protein were loaded on the gel. Molecular masses of proteins (given on the right) were judged according to a molecular mass standard run on the same gel. Staining with Ponceau S (the prominent band corresponding to the large subunit of RuBisCO) served to demonstrate equal loading. The experiment was performed in three independent biological replicates.

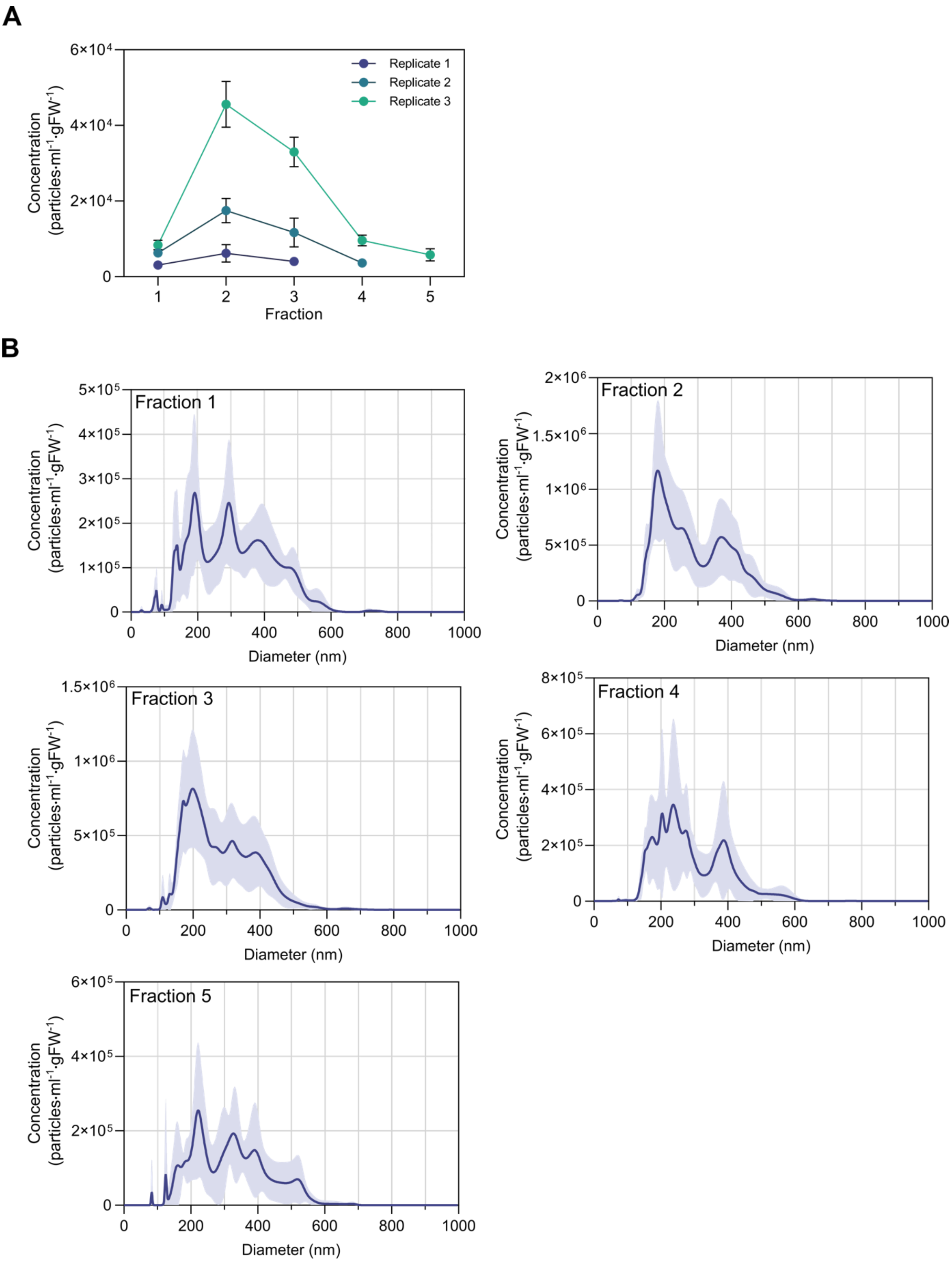


**Supplemental Figure 7. EVs elute in size exclusion chromatography fractions 2-4.**

**A** Mean particle concentration in SEC fractions 1-5 measured by NTA. Points represent the mean, error bars the standard deviation. EVs were isolated from leaves of 10-day-old barley that were sampled at 72 hpi with *B. hordei*. PME EV pellets were resuspended in 500 µl Tris (pH7.5)/1X resuspension buffer and subjected to SEC. Eleven to sixteen fractions were collected. Results of all fractions that contained enough particles for measurement are displayed. The experiment was performed in three independent biological replicates.

**B** NTA data showing the size distribution and mean concentration within the upper and lower 95% confidence intervals of PME-derived EV fractions. Plots are based on the dataset shown in panel A.

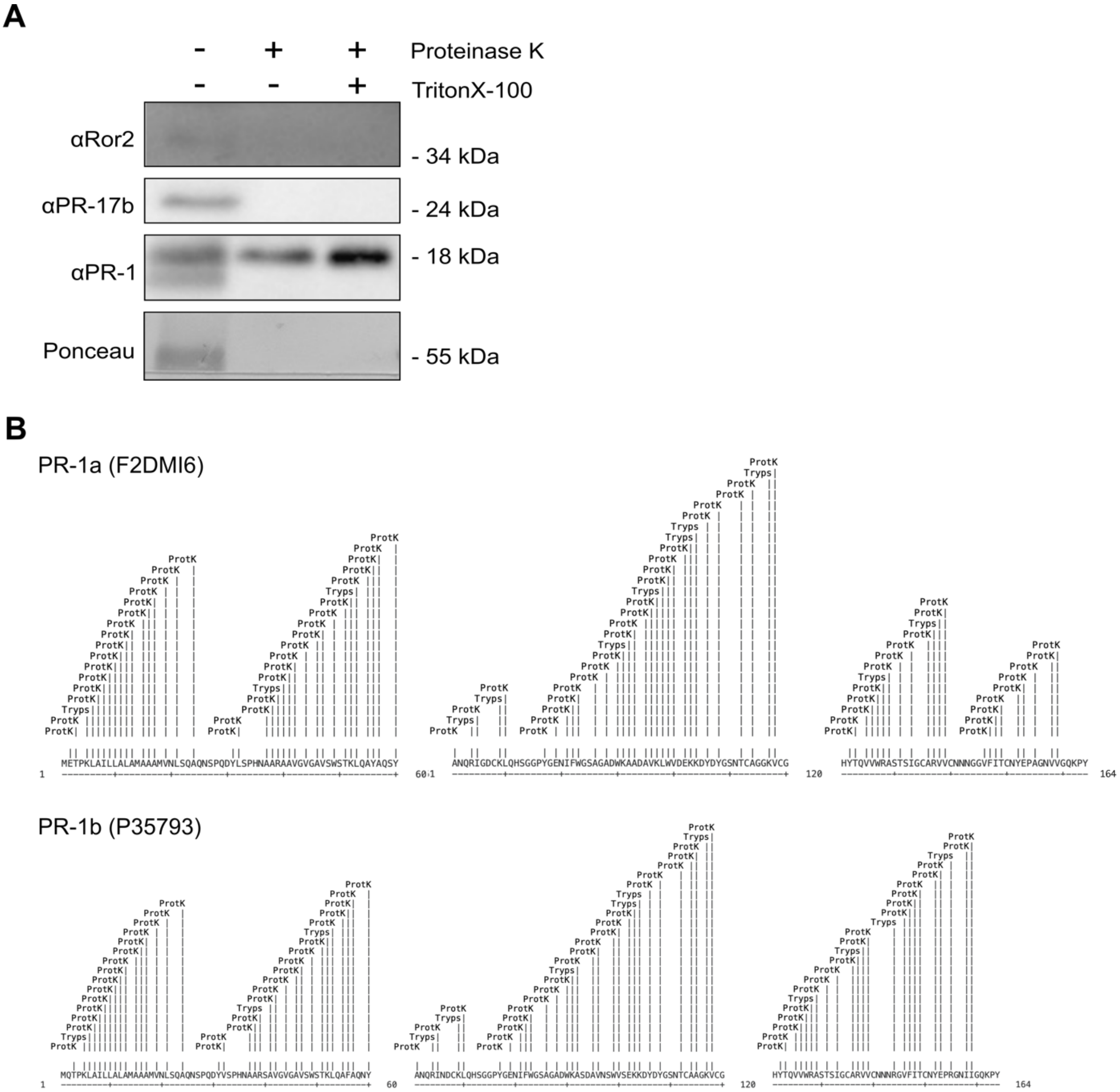


**Supplemental Figure 8. Predicted proteinase K and trypsin cleavage sites in PR-1a and PR-1b.**

**A** Immunoblots probed with antibodies specific for the potential EV marker protein Ror2 (predicted molecular mass 34 kDa), the defence protein PR-17b (predicted molecular mass 24 kDa), and the secreted defence marker PR-1 (predicted molecular mass 18 kDa) in PME-derived EVs in barley leaves sampled at 72 hpi with *B. hordei*. EVs were treated with either proteinase K or Triton X-100 or co-treated with Triton X-100 and proteinase K. Gels were loaded with 20 µl of treated EV pellets and mixed with 6x loading dye. Molecular masses of proteins (given on the right) were judged according to a molecular mass standard analysed on the same gel. The experiment was performed in four independent biological replicates.

**B** Proteinase K (ProtK) and trypsin (Tryps) cleavage sites in PR-1a (upper panel) and PR-1b (lower panel) were predicted with the Expasy Peptide Cutter (https://web.expasy.org/peptide_cutter/) using the respective UniProt amino acid sequences and standard settings.
